# Asian wild apples threatened by gene flow from domesticated apples and by their pestified pathogen

**DOI:** 10.1101/2020.04.15.042242

**Authors:** Alice Feurtey, Ellen Guitton, Marie De Gracia Coquerel, Ludovic Duvaux, Jason Shiller, Marie-Noëlle Bellanger, Pascale Expert, Mélanie Sannier, Valérie Caffier, Tatiana Giraud, Bruno Le Cam, Christophe Lemaire

## Abstract

Massive gene flow between crops and their wild relatives may threaten the genetic integrity of wild species. Such threats are now well documented, but little is known about indirect consequences involving the spillover of crop pathogens to wild plants or introgression between crop and wild pathogens. To address these questions, we used population genetics approaches, demographic inference and pathogenicity tests on host-pathogen pairs composed of wild or domesticated apple trees of Central Asia and their fungal pathogen, *Venturia inaequalis*, itself showing differentiated agricultural-type and wild-type populations. We confirmed the occurrence of gene flow from cultivated to wild apple trees in Asian forests, threatening the Asian wild apple genetic integrity. SNP markers and demographic modeling revealed the occurrence of a secondary contact followed by hybridization between agricultural-type and wild-type fungal pathogen populations, and the dispersal of the agricultural-type pathogen in wild forests. We detected a SNP predicting the ability of the fungus to parasitize the different host populations, which induced an early stop codon in a gene coding for a small secreted protein in the agricultural-type fungal population, thus representing a putative avirulence gene which function loss would enable to parasitize cultivated apples. Pathogenicity tests in fact revealed the pestification of *V. inaequalis*, with higher virulence of the agricultural-type population on both wild and domesticated trees. Our findings highlight the threat posed by cultivating a crop near its center of origin, with the invasion of a pestified pathogen on wild plants and introgression in the wild-type pathogen.

## Introduction

The domestication of plants corresponds to genetic and phenotypic differentiation between crops and their wild relatives under human selection (Zeder, Emshwiller, Smith & Bradley, 2006). However, crops and their wild relatives often remain interfertile, which can lead to introgression when the two taxa remain geographically close or come into secondary contact (Ellstrand et al., 2013). The hybrids resulting from crop-to-wild pollination events may have a low fitness in natural environments (Wang, Viera, Crawford, Chu & Nielsen., 2017), potentially driving to extinction the wild lineages receiving massive gene flow from its relative crop (Todesco et al., 2016; Wolf, Takebayashi & Rieseberg, 2001). In contrast, hybrids can outcompete pure wild individuals, thus threatening the genetic integrity of wild lineages (Feurtey, Cornille, Shykoff, Snirc & Giraud, 2017; Hooftman, Jong, Oostermeijer & Den Nijs, 2007; Hovick, Campbell, Snow & Whitney, 2012), thereby jeopardizing future adaptation to global changes and leading to a loss of valuable genetic resources for breeding programs.

A much less widely studied, but just as alarming, consequence of secondary contact between crops and their wild relatives, is the possible gene flow between their respective pathogens, which are themselves often closely related. Such gene flow between differentiated pathogen populations or species parasitizing wild and domesticated hosts can promote the emergence of new diseases or the breakdown of resistance, through the generation of pathogens with an expanded host range (Depotter, Seidl, Wood & Thomma, 2016), higher resistance to antibiotics (Hanage, Fraser, Tang, Connor & Corander, 2009) or enhanced virulence (i.e. degree of damage caused by the pathogen to its host) (Stukenbrock, Christiansen, Hansen, Dutheil & Schierup, 2012), thus potentially representing a major threat to both crop health and wild host persistence (Lemaire et al., 2016; Leroy, Lemaire, Dunemann & Le Cam, 2013; Leroy et al., 2016). Pathogen spillover (*i*.*e*., infection by a pathogen on another host than its endemic host) and hybridization between closely related phytopathogenic fungal species following secondary contact have been reported in several systems (Feurtey & Stukenbrock, 2018; Gladieux et al., 2011; Stukenbrock, 2016 a, b). Indeed, when fungal populations have diverged in allopatry and have specialized on new hosts and later met in sympatry (i.e., secondary contact), pre-mating barriers are often weak (Giraud, Refrégier, Le Gac, de Vienne & Hood, 2008; Le Gac & Giraud, 2008) and they can hybridize and/or still infect their ancestral host. The resulting spillover and hybridization events can lead to disease emergence and/or increased disease severity (Ioos, Andrieux, Marçais & Frey, 2006; Lin et al., 2007; Newcombe, Stirling, McDonald & Bradshaw, 2000; Stukenbrock et al., 2012). In anther-smut *Microbotryum* fungi for example, two sister castrating pathogens parasitize two sister plant species and co-occur in sympatry as a consequence of a secondary contact following allopatric divergence (Gladieux et al., 2011). One of the two species, *Microbotryum lychnidis-dioicae* (DC. ex Liro, Deml & Oberw.), is however better at infecting both hosts, and spillovers occur in nature (de Vienne, Hood & Giraud, 2009; Gladieux et al., 2011). In poplars, hybrid trees were generated and planted as they were resistant to the two rust species parasitizing the two parental tree species. However, a fungal hybrid between the two rust species rapidly emerged that was able to cause disease on the hybrid poplars (Newcombe et al., 2000). Yet, the consequences of crop-to-wild gene flow on the evolution of wild pathogen populations and on the disease of their wild host has been little investigated. Joint analyses of both wild and agricultural hosts and pathogens are required for a comprehensive understanding of the consequences of secondary contacts of crops and their wild relatives, and of their pathogens, as well as for elucidating coevolutionary histories and local adaptation. Yet, such studies combining both host and pathogen population genetic analyses remain scarce (Croll and Laine 2016).

Here, we addressed these questions on apple trees and their apple scab fungal pathogen, *Venturia inaequalis* (Cooke) G. Winter, which represent good models for investigating the consequences of secondary contact between crops and their wild relatives. They indeed have both wild and agricultural differentiated species/populations that have recently been reunified in their center of origin and may thus possibly hybridize (Figure 1). The cultivated apple (*Malus domestica* Borkh.) is the most cultivated fruit tree of temperate areas worldwide (FAO STATS from 2008-2018, as accessed on 04-07-2020). Analysis of phenotypic diversity, genetic markers, archeological data and demographic inference showed that the cultivated apple was initially domesticated from *M. sieversii* Ledeb. M. Roem (Figure 1A), forming quasi-monospecific natural forests of wild apple in the Tian Shan Mountains in Central Asia (Cornille et al., 2012; Harris, Robinson & Juniper, 2002; Vavilov, 1931; Velasco et al., 2010), which have been declared a World Heritage Center by UNESCO. The cultivated apple was imported into Europe by the Romans *via* the Caucasian and Mediterranean coasts, crossing the distribution ranges of various wild apple tree species on the way, leading to major secondary contributions to the cultivated apple tree genepool from several crabapples (Cornille et al., 2012; Cornille, Gladieux & Giraud 2013; Nikiforova, Cavalieri, Velasco & Goremykin, 2013). There has been in particular, in Western Europe, an important contribution from the European crabapple, *M. sylvestris* (L.) Mill, to the *M. domestica* genome (Figure 1A). Domestication and breeding has led to high differentiation between *M. domestica* and its various wild progenitors (Cornille et al., 2012). However, because apple species generally have weak interspecific barriers, introgression can still occur between wild and cultivated forms (Cornille et al., 2012; Nikiforova et al., 2013). This situation raises serious concerns regarding possible gene flow and its deleterious consequences for the conservation of important wild genetic resources. Yet, most studies on gene flow between domesticated and wild apples have focused on the European wild apple tree. These studies have shown that extensive crop-to-wild gene flow is threatening the genetic integrity of the European crabapple *M. sylvestris* (Cornille et al., 2015; Feurtey et al. 2017; Figure 1A). In Central Asia, at the end of the 19th century, western European *M. domestica* apple varieties have been planted in orchards close to *M. sieversii* forests, leading to recent secondary contact between *M. domestica* and *M. sieversii*. A couple of studies have suggested the occurrence of gene flow from *M. domestica* to the Asian crabapple *M. sieversii*, although without investigating the contact sites that orchards represent (Cornille et al. 2012; Omasheva et al., 2017). Such gene flow may affect the genetic integrity of *M. sieversii*, which would amplify the current threat on Asian wild-apple forest ecosystems, which are already endangered by forest destruction, *M. sieversii* being included in the International Union for Conservation of Nature’s Red List of Threatened Species (IUCN Red List) as a vulnerable organism.

**Figure 1:**
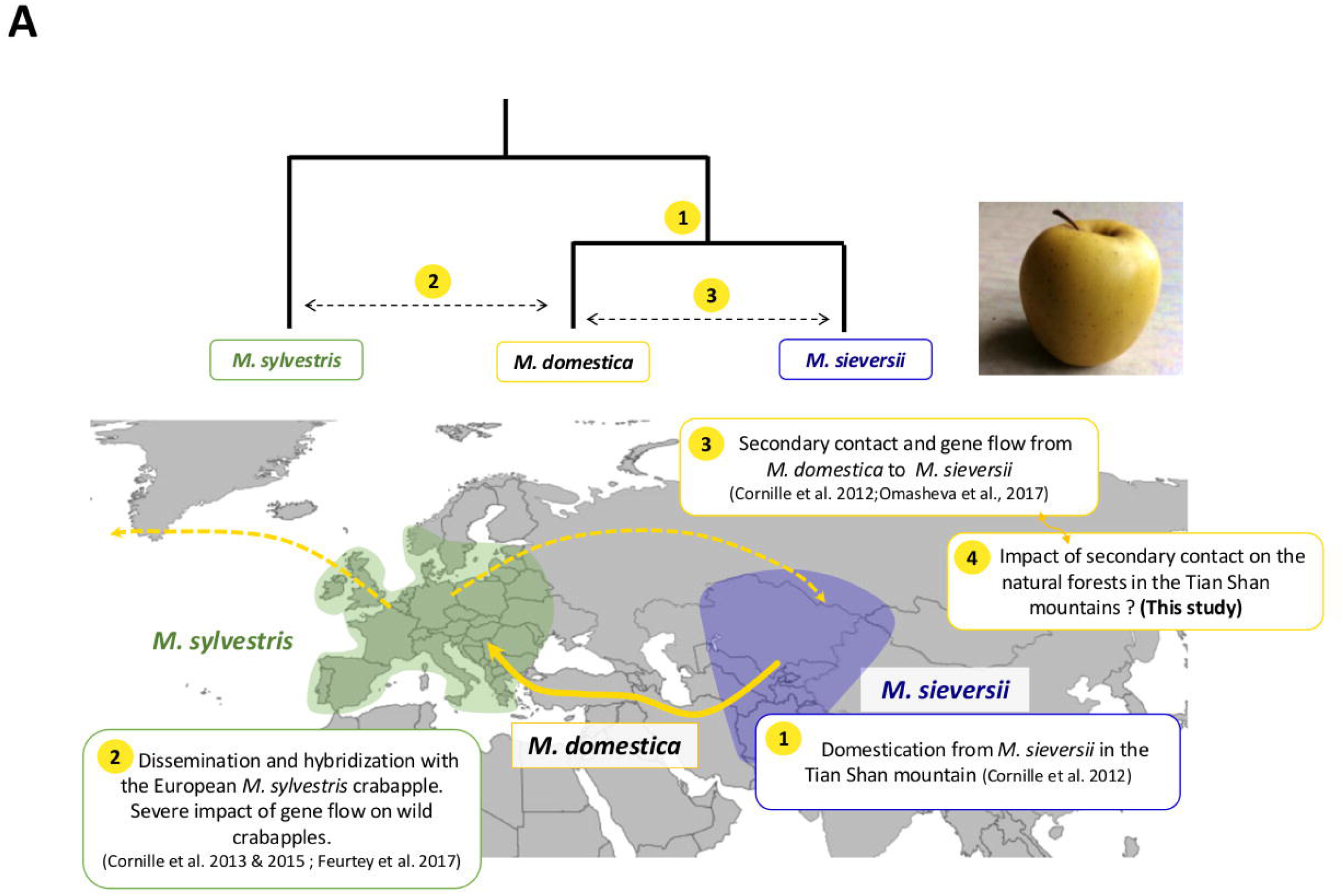
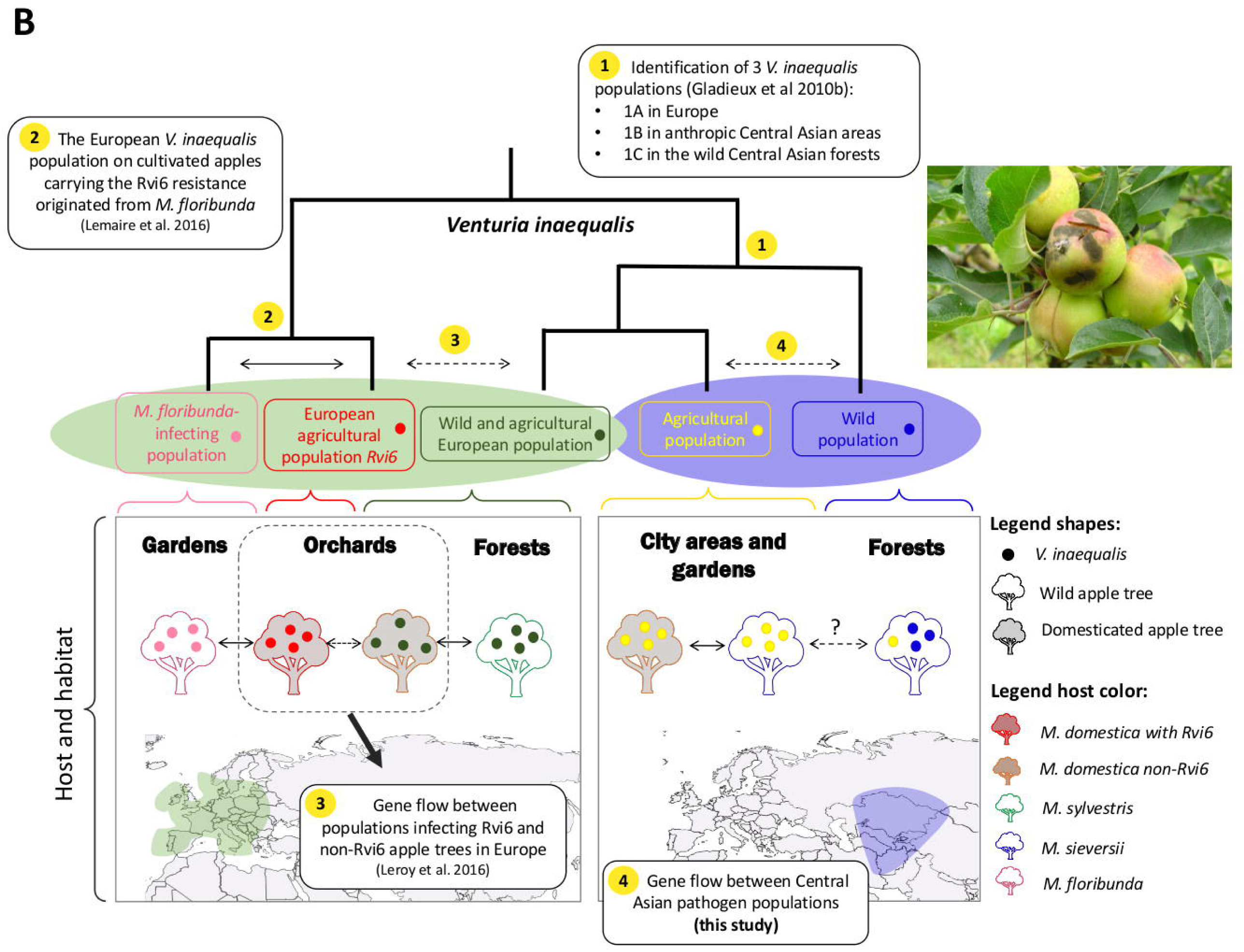
Summary of previous studies and questions addressed in the current study on *Malus* apple trees (A) and on their fungal pathogen *Venturia inaequalis* (B). **A**. The divergence and gene flow are shown at the top between the domesticated apple tree *M. domestica*, its initial progenitor *M. sieversii* from Asia and its secondary contributor the European crabapple tree *M. sylvestri*s (Cornille et al. 2012). The distribution areas of the two wild apple trees are shown at the bottom, indicating (1) the initial domestication in Asia (Cornille et al. 2012), migration along the Silk Road (yellow full line) and (2) secondary gene flow between *M. domestica* and *M. sylvestris* in Europe (Cornille et al. 2013; Feurtey et al. 2017), followed by dispersion of modern cultivars worldwide (dashed yellow arrows). (3) A couple of previous studies have investigated the consequence of the secondary contact between the domesticated apple and its Asian progenitor in terms of gene flow from *M. domestica* to *M. sieversii* (Cornille et al. 2012; 2012; Omasheva et al., 2017), and here we study the consequences of planting orchards in forest. **B**. The divergence is shown at the top between the fungal pathogen *V. inaequalis* populations and their distribution area at the bottom, with colored trees symbolizing apple tree species and colored circles the fungal populations. (1) Gladieux et al. (2010b) showed the existence of differentiated populations in Europe on *M. silvestris* and *M. domestica* (1A), in anthropic Central Asian areas on *M. sieversii* and *M. domestica* (1B) and in the wild Central Asian forests on *M. sieversii* (1C). (2) Lemaire et al. (2016) showed that the differentiated European *V. inaequalis* population on cultivated apples carrying the Rvi6 resistance originated from *M. floribunda*. (3) Leroy et al. (2016) investigated the existence of gene flow between populations infecting Riv6 and non-Riv6 apple trees in Europe. (4) The present study investigates the occurrence of gene flow among Central Asian pathogen populations, i.e., the population in anthropic Central Asian areas on *M. sieversii* and *M. domestica* and the population in the wild Central Asian forests on *M. sieversii*.

Apple domestication has also fostered divergence in the ascomycete fungus *V. inaequalis*, a haploid fungus with an obligatory sexual reproduction event each year. This fungus is responsible for the apple scab disease, producing gray-brown lesions on leaves and fruits, and leading to major economic losses on cultivated apples. *Venturia inaequalis* parasitizes *M. domestica* and several wild apple tree species, such as *M. sieversii, M. sylvestris* and *M. floribunda* (Siebold ex Van Houtte), and differentiated populations occur on these different *Malus* species (Figure 1B; Gladieux et al., 2010b). Apple trees and *V. inaequalis* share a common geographical origin in Central Asia (Cornille et al., 2012; Gladieux et al., 2008), *M. sieversii* being the wild host of origin of the fungus (Gladieux et al., 2010b). In the Central Asian Mountains, in forests where *M. domestica* is absent, a wild-type *V. inaequalis* population that represents a relic of the ancestral population has been found on *M. sieversii* trees (Gladieux et al., 2010b). An agricultural-type *V. inaequalis* population has been found in Central Asia (Gladieux et al., 2010b) in the peri-urban or agricultural environments on *M. domestica* and on *M. sieversii* (Figure 1B). These wild-type and agricultural-type populations of *V. inaequalis* began diverging in Central Asia between 2,000 and 4,000 years ago (Gladieux et al., 2010b), the agricultural-type population having then spread into Europe together with the domesticated apple (Figure 1B; Gladieux et al., 2010b). It has not been studied yet whether the current co-occurrence of the agricultural-type and wild-type *V. inaequalis* populations represents a secondary contact between the two fungal populations, and what are its epidemiological consequences through potential spillover and introgressions (Wang et al., 2017). Indeed, gene flow between wild-type and agricultural-type *V. inaequalis* populations may lead to the emergence of hybrids harboring new epidemiological traits, potentially harmful to both wild and cultivated apple trees, and/or to the dispersal of the agricultural-type population into wild forests. As a matter of fact, previous studies have shown that the *V. inaequalis* population parasitizing *M. floribunda*, and highly differentiated from all the populations on *M. sieversii* and *M. sylvestris*, migrated to resistant varieties of *M. domestica*, which promoted a resistance breakdown and gene flow between pathogen populations (Figure 1B; Lemaire et al. 2016; Leroy et al. 2016).

Despite the importance of wild Asian apple trees as an endangered wild species and as a valuable genetic resource for future breeding programs, the potential impacts of the secondary contact between domesticated and wild apple trees in Central Asia on the fitness of both the plant and its associated pathogen have not been investigated yet. The lack of diagnostic morphological features for apple species makes the use of genetic markers necessary for species and hybrid identification (Cornille, Giraud, Smulders, Roldán-Ruiz & Gladieux, 2014). We therefore used microsatellite markers in apple trees and genome-wide single-nucleotide polymorphisms (SNPs) in *V. inaequalis*, together with pathogenicity assays, to assess the occurrence and impact of secondary contact on both fruit trees and their fungal pathogens in their center of diversity. First, we investigated whether the introduction of domesticated apple trees in Central Asia threatened the genetic integrity of *M. sieversii* wild apple trees via introgressions, and whether orchards planted in forests represent particular threats. We then assessed whether the agricultural-type fungal population occurred on wild apple trees and hybridized with the wild-type population. We tested, using demographic modeling and scenarios comparison, the hypothesis that the co-occurrence and hybridization of agricultural-type and wild-type *V. inaequalis* populations resulted from a secondary contact at the time when domesticated trees were introduced in orchards near wild apple forests, after a period of allopatry. We also tested whether some genomic regions in *V. inaequalis* were good predictors of its ability to parasitize the different host populations. Finally, we tested, using experimental inoculations, whether such gene flow increased the threats to the wild apple tree through increase in virulence of the pathogen populations.

## Material and Methods

### Sampling

Plant and fungal materials were sampled in 2012 at nine sites in Kazakhstan in the Tian Shan Mountains, the origin center of the apple tree domestication where the natural forest is dominated by wild apple trees (Cornille et al., 2014). Seven of the nine sampling sites were located in natural apple forests, farther than dozens of kilometers from any orchard. The two remaining sampling sites corresponded to a 20-30 year-old orchard and the wild natural forest immediately surrounding the orchard, respectively. We sampled apple trees in the orchard site and wild apple trees in the surrounding forest. This orchard was situated directly in the middle of the natural forest and constituted a rare situation allowing to investigate the consequences of the secondary contact for both the host and the pathogen. We obtained plant material from 245 apple trees, originating from the nine sites. The 245 plant samples corresponded to 185 leaves and 60 fruit skin pieces, all harbouring visible scab lesions (Table S1). No morphological differences between *M. domestica* and *M. sieversii* allow to distinguish them reliably in the field. Therefore, all sampled trees were genotyped with 33 microsatellite markers and were assigned to *M. domestica* or *M. sieversii* species or to a hybrid class using as references 50 apple genotypes previously characterized as being non-hybrid *M. domestica* genotypes and 25 apple genotypes previously characterized as being non-hybrid *M. sieversii* genotypes (Cornille et al., 2012).

Monospore isolation of the fungus was performed from disease lesions on 205 of these 245 trees: 24 to 48 h after spreading a spore suspension obtained from the diseased material on a Petri dish with malt-agar medium, we took a single germinated spore using a needle under a stereomicroscope for culture on a new Petri dish. We performed one monospore isolation per tree for 148 trees and two to four monospore isolations per tree for 57 trees. In total, we obtained 269 strains (Table S1). In addition to the fungal collection sampled for this study, we included in our final dataset 15 *V. inaequalis* strains previously assigned to the pure wild type population and 18 *V. inaequalis* strains previously assigned to the agricultural-type population (Gladieux et al., 2010b). These 33 *V. inaequalis* reference strains whose genomes have been previously sequenced (Le Cam et al., 2019) had been sampled in Kazakhstan on *M. sieversii*, either from a peri-urban environment near Almaty (43°15000N-76°54000E) or from a natural forest site in the Tian Shan Mountains (43°13807N-77°16783E). Despite being sampled on *M. sieversii*, the 18 strains from the peri-urban environment clustered with strains sampled on domesticated apples in Kazakh orchards, thus belonging to the agricultural type (Gladieux et al., 2010b) (Figure 1B; Table S2).

### SNP design and genotyping for the apple scab fungus *Venturia inaequalis*

We used the set of single nucleotide polymorphisms (SNPs) previously detected in the genomes of the 15 wild-type and 18 agricultural-type reference strains (Gladieux et al., 2010b; Le Cam et al., 2019). SNPs were filtered in order to avoid transposable elements and AT-rich regions, as well as more than 20% of missing data (Le Cam et al., 2019). After filtering, the dataset contained 1,210,121 SNPs, with no bias of missing data frequencies between wild and agricultural populations. In order to identify a set of SNPs distinguishing agricultural and wild-type *V. inaequalis* strains, we selected 192 SNPs among the 1,210,121 available SNPs as those meeting three requirements: *i)* with moderate (*F*_*ST*_ = 0.21) to high (*F*_*ST*_ = 1) differentiation between wild-type and agricultural-type reference populations (Figure S1, Table S3), to obtain reasonable statistical power to assign genotypes to wild, agricultural and hybrid types, *ii)* physically distant enough to ensure lack of linkage disequilibrium between them (correlation coefficient between markers within population r^2^ < 0.1), and iii) found in predicted genes (Le Cam et al., 2019).

For each monospore strain, DNA was extracted from 30 to 40 mg of mycelium as previously described (Leroy et al., 2016). The 269 strains were genotyped at the 192 SNP markers (Table S3) using the *KASPAR* method (KBioscience, competitive allele-specific polymerase chain reaction assay). *KASPAR* genotyping was performed at the *Gentyane* platform (INRA, Clermont-Ferrand, France) using 10 ng of haploid DNA mixed with the KASP genotyping master mix (Catalogue number: KBS-1016-017, LGC Genomics, Hoddesdon, UK) and custom KASP SNP assays (LGC Genomics, Hoddesdon, UK). Only SNPs and individuals with less than 20% missing data were kept for analyses. After filtering for missing data, the dataset was composed of 255 fungal strains genotyped for 181 SNPs. Details about SNP location on the *V. inaequalis* reference genome are given in Table S3.

### Genotyping using microsatellite markers for the host apple tree

In order to assign trees to wild, domesticated and hybrid types, all apple trees were genotyped using microsatellite markers. Apple tree DNA was extracted from leaves with the NucleoSpin plant DNA extraction kit II® (Macherey & Nagel). Microsatellite PCR amplifications were performed with a Multiplex PCR Kit® (QIAGEN, Inc.). To genotype the 245 trees, we used 33 microsatellite markers spread across the 17 chromosomes using 10 different multiplex reactions, as previously described (Cornille et al., 2012). The genotyping was done at the *Gentyane* platform (INRA, Clermont-Ferrand, France). Only the microsatellite markers with less than 30% of missing data were used and only individuals with less than 50% of missing data were kept for the analyses. The final dataset was thus composed of 240 apple tree samples genotyped at 28 microsatellite markers.

Since fungi were sampled on the genotyped host trees, we had genotypes for host and pathogen pairs. We obtained genotypes for 249 host-pathogen pairs, several fungal samples originating from the same trees (132 trees with one fungal strain, 49 trees with two strains, five trees with three strains and one tree with four strains).

### Analyses of diversity, differentiation and hybridization

Reference genotypes of *M. domestica* (N=50) and *M. sieversii* (N=25) (Cornille et al., 2012) were used to estimate the proportion of ancestry from these two species in the 240 genotyped apple trees. We computed a hybrid index (*HI*) implemented in the add-on R-package *Introgress* (Gompert & Buerkle, 2010), an estimate of the proportion of alleles that were inherited from one of the two parental populations, *i*.*e*. the level of ancestry in one of two populations. In our case, we set pure *M. domestica* trees to *HI*=1 while *M. sieversii* had *HI*=0. This index therefore represents the proportion of ancestry in the *M. domestica* genepool (thus called here *P*_*dom*_). For the pathogen, we used the same package to compute a hybrid index based on the reference genotypes of strains previously identified as belonging to the agricultural-type population (N=18) or to the wild-type (N=15) population (Gladieux et al., 2010b). Here, we set *HI* = 1 for the agricultural type, so that the hybrid index represents the proportion of ancestry in the agricultural-type population (here called *P*_*agr*_) genepools.

In most of our analyses, we used the hybrid indices as continuous values, and not discrete classes, to avoid potential biases generated by arbitrary thresholds. Only for the analyses of apple and fungal population or species spatial distribution, of the genetic determinants of adaptation and of pathogenicity tests, we assigned apple and fungal genotypes to discrete classes. Apple tree genotypes with a *P*_*dom*_ index below 0.2 and those with a *P*_*dom*_ index greater than 0.8 were considered to belong to *M. sieversii* and *M. domestica*, respectively, based on the distribution of the hybrid index (Figure 2A). A fungal strain with a *P*_*agr*_ below 0.1 was considered to be a wild-type strain, whereas a strain with a *P*_*agr*_ greater than 0.9 was considered to be an agricultural-type strain, based on the distribution of the hybrid index (Figure 2B). The threshold was more stringent for fungi than for trees, as more markers were available and the power was, therefore, higher. For both pathogens and hosts, genotypes with intermediate hybrid index values were considered to be hybrids. Assigning hybrids to more precise hybrid classes (*ie* F1, F2 or backcrosses) is not possible with the current dataset for haploid organisms such as *V. inaequalis*. When we used discrete classes for analyses, we checked that the choice of the thresholds did not affect the inference.

**Figure 2:**
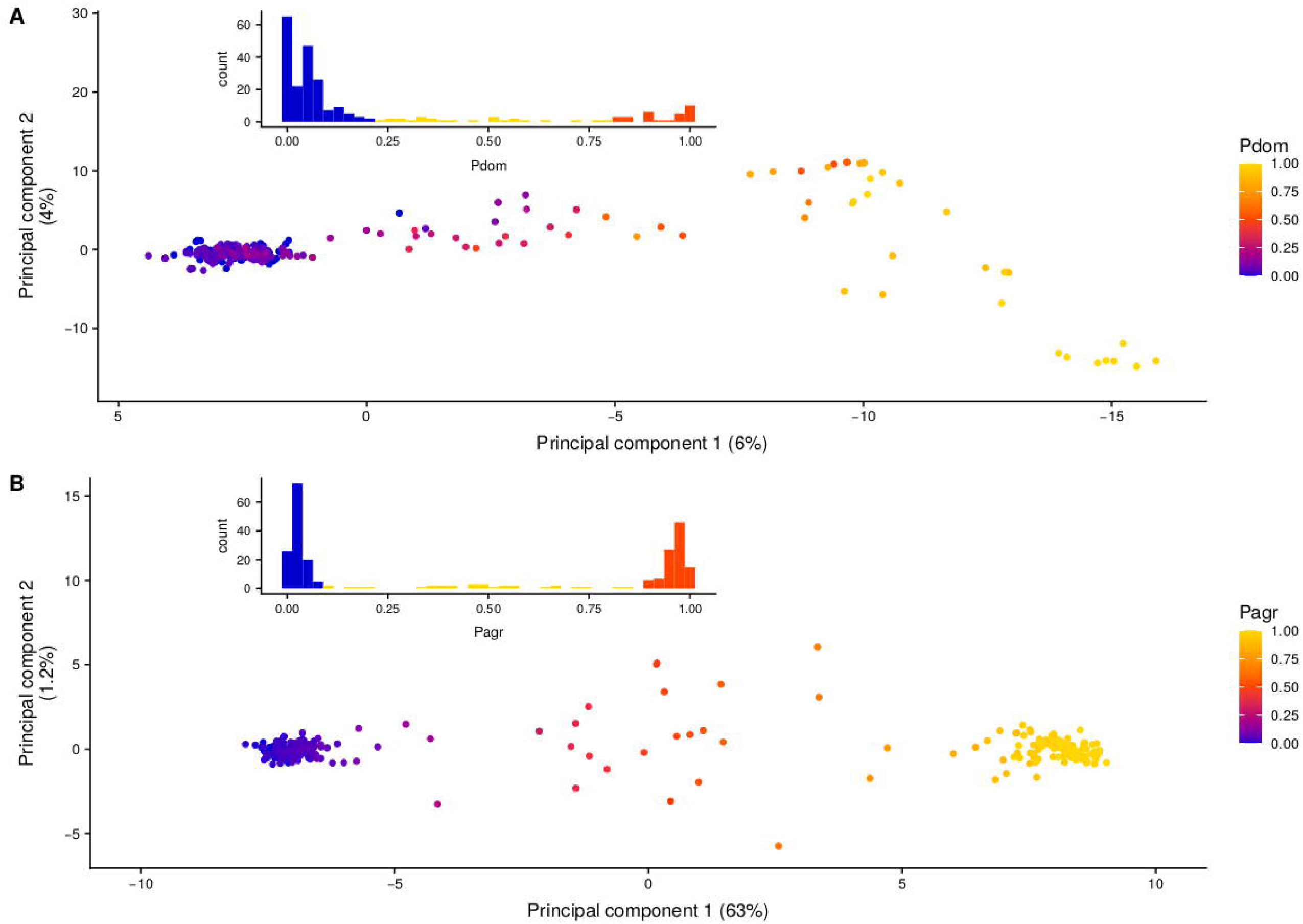
Genetic diversities and ancestry assignments for apple trees and *Venturia inaequalis*. (A) Principal component analysis on genotypes of 240 apple trees (*Malus spp*.) for 28 microsatellite markers, with their ancestry assignment (*P*_*dom*_) to the cultivated apple tree *M. domestica* indicated by color. The distribution of the *P*_*dom*_ index is shown in the inset at the top left. (B) Principal component analysis on genotypes of 255 *V. inaequalis* strains for 181 SNPs, with their ancestry assignment (*P*_*agr*_) to the agricultural-type of *V. inaequalis* indicated by color. The distribution of the *P*_*agr*_ index is shown in the inset at the top left.

For both apple and *V. inaequalis* samples, we also investigated genetic differentiation and species or genetic lineage assignment through principal component analyses (PCAs) using the ade4 R package (Dray & Dufour, 2007) based on the multilocus genotypes (microsatellite markers for trees and SNPs for fungi). We further investigated the existence of population structure within apple tree species and within the previously delimited *V. inaequalis* populations using the software STRUCTURE v.4.3 (Falush, Stephens & Pritchard, 2003) based on 10 runs of 250,000 Monte Carlo Markov Chain (MCMC) iterations after a burn-in of 25,000 iterations using the admixture model and assuming a number of clusters ranging from 1 to 6. The output of STRUCTURE was processed with Clumpp (Jakobsson & Rosenberg, 2007) and we used the method described in Evanno et al. (2005) and implemented in STRUCTURE HARVESTER (Earl & vonHoldt, 2012) to detect the strongest population subdivision level. We tested the significance of correlations between the hybrid degrees inferred from coordinate values of the first PCA axis, the hybrid index and the ancestry probability given by STRUCTURE using a Pearson’s product-moment correlation coefficient in R.

For both apple and *V. inaequalis* populations, we estimated classical population genetics statistics, i.e., expected heterozygosity (*H*_*e*_, Weir & Cockerham, 1984), as well as global and pairwise differentiation *F*_*ST*_ (Weir & Cockerham, 1984) between the nine sampling sites. All statistics were computed and tested for significance using GENEPOP v.4.7.5 (Rousset, 2008). Significativity of global and pairwise *F*_*ST*_ was assessed using Fisher’s exact tests of allele frequency differences between sampling sites with 10,000 dememorization and batches and 10,000 iterations per batch. We also performed Mantel tests of correlation (Mantel, 1967) between pairwise *F*_*ST*_ matrices computed for apple and *V. inaequalis* populations on the one hand, and between each pairwise *F*_*ST*_ matrix and a matrix of geographical distance between sampling sites on other hand, using the *mantel*.*test* function of the ape v5.3 R package (Paradis & Schliep, 2019).

### Genetic footprints of host adaptation in *Venturia inaequalis*

In order to detect genetic signatures of adaptation to host species in *V. inaequalis* populations, we performed a discriminant analysis on principal components (DAPC; Jombart, Devillard & Balloux, 2010) implemented in the R package *adegenet v*.*2*.*12* (Jombart, 2008). Unlike PCA which focuses on overall genetic variance, this procedure seeks variables, the discriminant functions, that maximize differences between *a priori* delimited groups, while minimizing the variance within these groups. We applied this analysis by defining groups in *V. inaequalis* based on the assignment of their apple tree groups of collection (*M. domestica, M. sieversii* and hybrids). This approach also aims at detecting the genetic factors (here SNPs) that contributed the most to this ecologically-based clustering. If one of the 181 SNPs in *V. inaequalis* was closely linked to a genetic variant involved in adaptation to host species, its loading on the first discriminant functions would indeed be greater than that of other SNPs. In addition, we estimated monolocus pairwise *F*_*ST*_ between the three host-based fungal clusters, using Genepop v4.7.5 (Rousset, 2008).

To assess what allele was ancestral or derived at the SNP found to predict the host species of collection, we used five available genomes (Le Cam et al., 2019) belonging to the outgroup *V. inaequalis* lineage parasitising Pyracantha (Gladieux, Caffier, Devaux & Le Cam, 2010a; Le Cam, Parisi & Arene, 2002), with the following NCBI accession numbers SAMN07816619, SAMN07816620, SAMN07816621, SAMN07816617, SAMN07816618, corresponding to the strains 186, 1669, 2266, 2269 and 2507 respectively (Le Cam et al., 2019).

### Pathogenicity tests of *V. inaequalis* on *M. domestica* and *M. sieversii*

In order to test for an effect of the agricultural, hybrid or wild-type status of *V. inaequalis* strains on their pathogenicity on *M. domestica* and *M. sieversii*, we performed artificial inoculations using 57 strains from our sampling, corresponding to 20 strains identified as agricultural-type, 17 strains identified as wild-type and 20 hybrid strains (Table S1). We used one variety of *M. domestica* (GALA^®^, X4712) and one accession of *M. sieversii* (GMAL 3619.b) for the pathogenicity tests. Gala is a variety planted worldwide despite being highly susceptible to apple scab. GMAL 3619.b has been collected from the Tarbagatai mountain range by American apple breeders (Forsline, Aldwinckle, Dickson, Luby & Hokanson, 2003) and then introduced in France. We grafted the plants on the rootstock MM106 to obtain a vigorous growth favorable to disease development and we inoculated *V. inaequalis* strains when the plants were actively growing. Prior to inoculation, grafted plants were transferred to a quarantine-controlled climatic chamber. Because of the Asian origin of the strains, we used a quarantine chamber to prevent the risk of pathogen escape. Due to the lack of space in climatic chambers, we had to perform experiments at two different dates. In the first experiment, we used 10 agricultural-type strains, 10 hybrids and 8 wild-type strains. In the second experiment, we used 10 agricultural-type strains, 10 hybrids and 9 wild-type strains. In addition, for normalizing the two experiments, six strains (two strains of each type) that were used in the first experiment were also included in the second experiment and used as calibration strains in the analysis.

Each strain of *V. inaequalis* was grown on a cellophane sheet placed on malt-agar medium at 17°C to obtain spores (Caffier et al., 2014). The cellophane sheets were dried and stored at - 20°C. A spore suspension was made by collecting spores from these cellophane sheets and diluting them in water to a final concentration of 1.5×10^5^ spores mL^-1^ (Lê Van et al., 2012). For each strain, the spore suspension was inoculated on three Gala trees and three GMAL 3619.b trees using a mechanical air pressure sprayer. As trees actively grew during the experiment, we labelled the youngest fully deployed leaf of each tree as the F0 leaf, one or two days prior to inoculation to facilitate later disease scoring. To favor spore germination and fungal infection, the plants were kept in darkness with moisture maintained at 100% and temperature at 17°C for 48 hours after inoculation (Lê Van et al., 2012). Afterward, moisture was reduced to 80% during the day and 90% at night, with 12 hours of light per day. These climatic conditions are highly favourable to scab infection and thus provide a good indication of the strain ability to infect tree genotypes (MacHardy, 1996). For each host genotype, plants were randomized within three blocks to have in each block one host replicate for each strain. Disease severity, *i*.*e*., virulence of the fungal strain, was measured when there was no more increase of the disease symptoms: at 19 days post inoculation (dpi) for *M. sieversii* and at 21 dpi for *M. domestica*. Disease severity was measured as the percentage of a leaf displaying sporulation, from 0% when there was no disease symptom to 100% when the whole leaf was covered with spores. The F0 leaf being the last leaf deployed one or two days prior to inoculation, the assessment was done on the F0 leaf and on the first leaf under F0, named F1.

### Statistical analysis of phenotypic data

The data of the two distinct inoculation experiments were standardized based on the data from the six calibration strains: the disease severity values were corrected to obtain the same mean between the two distinct experiments for the six calibration strains. The virulence of each strain was estimated as the mean percentage of scabbed leaf area across the two leaves F0 and F1 and across the three inoculated plants of a given genotype (either *M. domestica* or *M. sieversii*), so that there was one value per fungal strain and per apple tree genotype in the analysis. As the assumptions of normality were not met, we compared the medians of virulence between the three different populations (agricultural-type, wild-type and hybrids) using a Kruskal-Wallis and a *post-hoc* Wilcoxon test at 21 dpi on *M. domestica* and 19 dpi on *M. sieversii* in R (version 3.4.4; R Core Team, 2018), considering the different strains in each population as replicates for the population effect. In addition, a Kendall test was performed to analyse the correlation between virulence and hybrid index *P*_*agr*_ for the 20 hybrid strains.

### Demographic inference for wild and agricultural types of the apple scab fungus *Venturia inaequalis*

In order to test demographic scenarios, and in particular the likelihood of divergence history without gene flow followed by a secondary contact in the fungal pathogen, we used the composite-likelihood, diffusion approximation-based approach for demographic inference implemented in ∂a∂i (Gutenkunst, Hernandez, Williamson & Bustamante, 2009). We used the 33 reference *V. inaequalis* genomes described above, belonging to 15 wild-type and 18 agricultural-type strains. In order to meet the requirement of marker independence, we thinned the 1,210,121 available SNPs for these genomes to keep only one SNP every 5kb (a threshold set based on the LD decay curve, Figure S2) using vcftools (Danecek et al., 2011), which left 6,187 SNPs. Using ∂a∂i (Gutenkunst et al., 2009), we tested a set of four basic models (Tine et al., 2014): strict isolation (SI) in which the two lineages diverge without gene flow, isolation with migration (IM) in which the two lineages diverge with constant gene flow, ancient migration (AM) in which the two lineages diverge with gene flow and stop exchanging genes at a time noted T_AM_, and secondary contact (SC) in which the two lineages diverge without gene flow and then exchange gene since a secondary contact occurring at T_SC_. Each of these four models was evaluated and fitted with the observed joint allele frequency spectrum (jAFS) using 20 independent runs. Each run started with perturbed starting parameters, from which a global optimisation was done with a simulated annealing optimisation procedure and was followed by an optimisation phase with a maximum number of 100 iterations (https://popgensealab.wordpress.com/dadi-inference/; Christe et al., 2017; Tine et al., 2014). For nested models, likelihood ratio tests (LRTs) were used to identify the best model. For non-nested models, the relative likelihood of Akaike criterion (AIC) was used instead (Christe et al., 2017).

Demographic parameters were estimated for the four models. These parameters included migration rates in the two directions, effective sizes in the two lineages, divergence time and time elapsed since other demographic events (T_AM_ in AM model and T_SC_ in SC model). The ancestral effective size before the split was calculated as *N*_*ref*_ = θ / (2 x *µ* x *L*) (Gutenkunst et al., 2009), where θ was estimated by∂a∂i; *µ* is the mutation rate per nucleotide per generation, estimated to be 2 × 10^−8^ from an analysis of divergence with the closely related species *Venturia pirina* parasitizing pear trees (Le Cam et al., 2019; data not shown). *L* is the effective genome length analyzed, being computed as G × s/S (Gutenkunst et al., 2009), where *G* is the size of the genome used, *S* is the total number of SNPs called in *G*, and *s* the actual number of SNPs used for inference. In our analysis, 1,210,121 SNPs were called on a portion of 34,942 Mb of the genome, 6,187 SNPs were used, so that *L* was computed as 34,942,000 × (6,187 / 1,210,121) = 178,648 bp. All other parameters were scaled by *N*_*ref*_. Times in years were computed using a generation time of one year, as *V. inaequalis* undergoes an obligate sexual reproduction once a year. The model parameters estimated and their 95% confidence intervals were obtained with Godambe methods (Coffman, Hsieh, Gravel & Gutenkunst, 2015) from 1000 bootstraps across SNPs.

## Results

### Contrasting distributions of the wild and agricultural host and pathogen populations in the Tian Shan Mountains

We explored the genetic makeup of apple trees and their scab pathogens in a wild environment, by sampling apple trees and fungal strains from the same trees, at nine sites in the forests of the Tian Shan Mountains in Kazakhstan. For apple trees, the PCA (Figure 2A) and STRUCTURE analysis (Figures S3A and B) confirmed the existence of two genetically distinct clusters corresponding to the orchard trees on the one hand, assigned to *M. domestica*, and most of the wild forest trees on the other hand, assigned to *M. sieversii*. We did not detect further population subdivision within the wild species (Figures 2A and S3). Both the first axis of the PCA and STRUCTURE analyses revealed footprints of admixture between the two tree species. The *P*_*dom*_ hybrid index values were also estimated for the 240 successfully genotyped trees. All three methods gave congruent results (Figs. 2A and S3C). The correlation coefficient between *P*_*dom*_ and the coordinates of the first PCA axis was high and significant (r=-0.97; p-value < 0.001; Figure S4A). The correlations between the ancestry proportion inferred by STRUCTURE and *P*_*dom*_ on the one hand, and between the ancestry proportion inferred by STRUCTURE and the coordinates of the first axis of the PCA on the other hand, were also high and significant (r=0.96, p-value<0.001, and r=-0.97, p-value<0.001, respectively). Based on the P_dom_ hybrid index, we identified 78% pure wild *M. sieversii* trees, ca. 12% *M. domestica* trees and near 10% hybrid trees (Figure 2A; Table S1). The species identified genetically were consistent with our expectations during sampling: only one *M. domestica* was identified in the natural forest sites (Figure 3), whereas most of the trees sampled in the orchard belonged to *M. domestica* (28 out of 30 trees sampled in the orchard, with 14 different genotypes). Three of the forest sites contained only *M. sieversii* trees, but the four other forest sites contained both *M. sieversii* and hybrid trees (with hybrid trees representing 11.1 to 27.6%, Figures 3), as did the forest site directly surrounding the orchard (with hybrid trees reaching there 27.3%).

**Figure 3:**
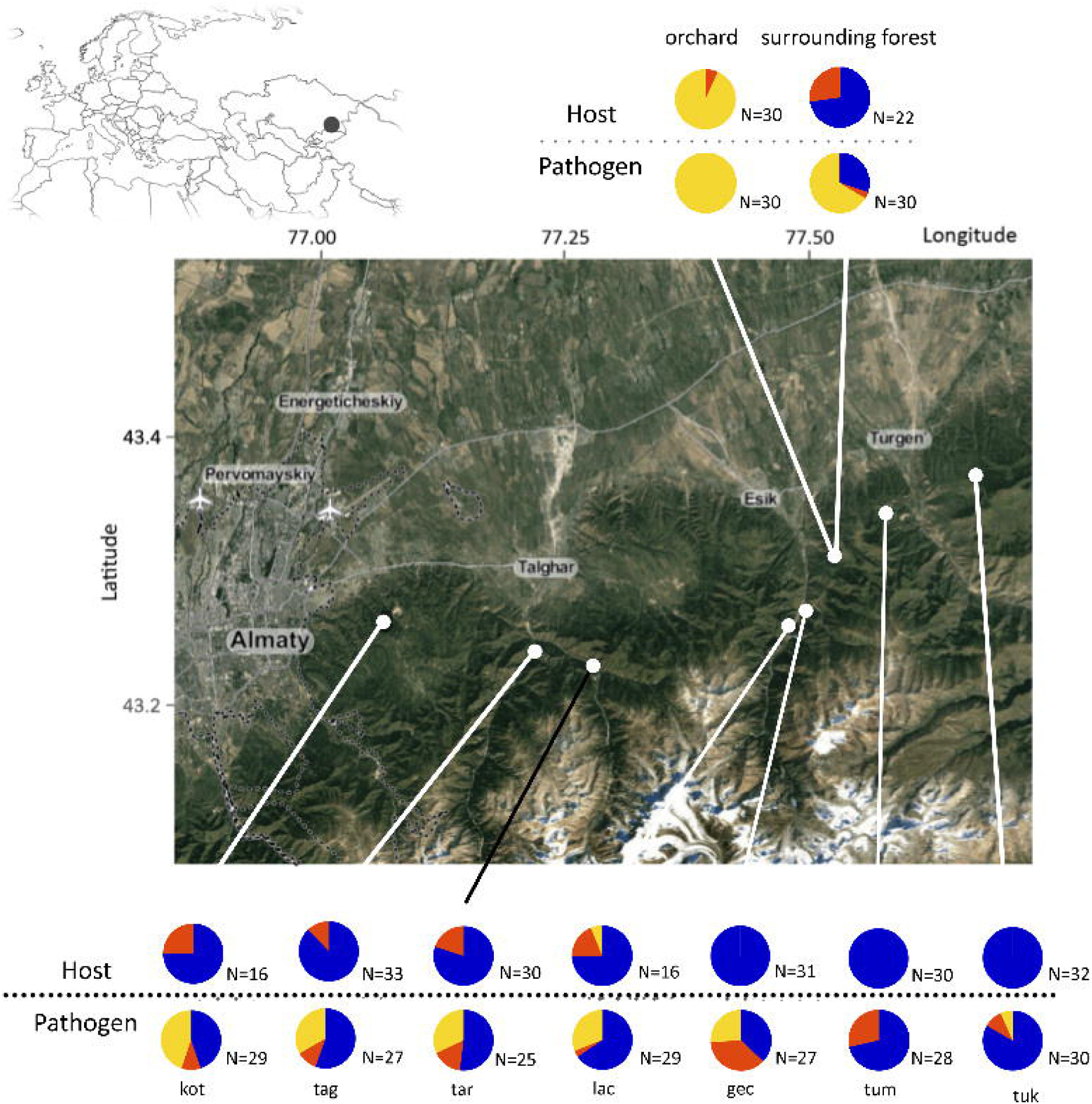
Distribution of the samples of apple trees and their fungal pathogen *Venturia inaequalis* in nine sites in the Tian Shan Mountains near Almaty (Kazakhstan). The black dot on the upper left corner map represents the global sampling location. Proportions of wild (blue), domesticated/agricultural (yellow) and hybrid (red) genotypes are shown for each sampling site for *Malus* trees (upper row of pie charts) and for their scab fungal pathogen (lower row of pie charts). The pie charts below the figure correspond to sampling sites in the natural forest. The pie charts above the figure refer to a sampling location composed of two sites, an orchard and the natural forest immediately surrounding it. Codes of the sampling locations: kot at Koturbulac; tag and tar at Talghar; lac, gec and esi (esi_o for the orchard and esi_f for the surrounding forest) at Esik; tuk and tum at Turgen.

For *V. inaequalis*, the PCA (Figure 2B) and STRUCTURE analysis (Figures S5A and B) confirmed the existence of two differentiated populations and the lack of further subdivision, and revealed admixture between the wild and agricultural populations (Figures 2B and S5B). The hybrid index *P*_*agr*_, the ancestry coefficient given by STRUCTURE and the first axis of the PCA, separating individuals according to their degree of ancestry into the two divergent populations (Figure 2B), gave congruent results (Figure S5C). The correlation coefficient between P_agr_ and the first axis of the PCA was high and significant (r=0.99; p-value=0.001) (Figure S4B). The correlations between the ancestry proportion inferred by STRUCTURE and *P*_*agr*_ on the one hand, and between the ancestry proportion inferred by STRUCTURE and the coordinates of the first axis of the PCA on the other hand, were also high and significant (r=0.99, p-value<0.001 and r=0.98, p-value<0.001, respectively).

The *P*_*agr*_ hybrid index estimated for the 255 *V. inaequalis* strains displayed a U-shaped distribution, with 33 fungal strains (12.9%) having *P*_*agr*_ values between 0.1 and 0.9, thus being considered to be hybrids, and with high frequencies of pure agricultural-type (38.5%) and pure wild-type (48.6%) fungal genotypes (Figure 2B; Table S1). All the fungal strains sampled in the orchard carried alleles typical of the agricultural-type (Figure 3), and the fungal strains sampled on the surrounding forest were mostly either agricultural-type or wild-type genotypes. A single hybrid strain was detected in the forest surrounding the orchard, whereas 3.4% to 37% of the fungal strains were hybrids in the other forest sites (Figure 3).

In order to further explore the relationship between host and pathogen genotypes, we compared their genetic diversity and differentiation across the sampling sites. Genetic diversities per sampling site ranged from 0.703 (tuk) to 0.802 (kot) for apple trees and from 0.244 (tuk) to 0.487 (kot) for *V. inaequalis* (Table S4). Genetic diversities were minimal and maximal in the same sampling sites for both the host and the pathogen, but no significant correlation was observed between the apple and fungus genetic diversity levels across sites (Kendall’s tau=0.389, p-value=0.18; Figure S6). Global *F*_*ST*_ among sites were 0.259 for apple trees and 0.243 for *V. inaequalis*. Pairwise *F*_*ST*_ estimates ranged from 0.000 to 0.207 for apple trees (Table S5) and from 0.000 to 0.721 for *V. inaequalis* (Table S6). For both the host and the pathogen, the orchard (esi_o) was the most differentiated from other sampling sites (Tables S5 and S6). Host and pathogen pairwise differentiation matrices were significantly correlated (Mantel-test Z-score=0.681, p-value=0.011; Figure S7). Yet, we found no significant correlation between pairwise *F*_*ST*_ and geographical distances matrices, neither for apple trees (Mantel test Z-score= 33.10, p-value=0.42) nor for *V. inaequalis* (Mantel test Z-score=103.20, p-value=0.51), indicating a lack of isolation by distance.

### All pathogen types occur on wild apple trees, but wild-type pathogens occur only on trees with no domesticated ancestry

We investigated the association between the fungal pathogen types and the *Malus* genotypes. From the 240 trees and 255 fungal strains successfully genotyped, we could obtain 249 host-pathogen pairs, several strains having been isolated from the same host trees. A sharp L-shaped pattern was observed when the hybrid index values of the host trees were plotted against those of their associated fungal genotype (Figure 4A). Hybrids were found among both trees and fungi, but no hybrid pathogens were collected from hybrid trees. Hybrid and wild-type fungi were found only on wild apple trees, whereas agricultural-type fungi were found on all types of apple trees, across the whole range of the hybrid index.

**Figure 4:**
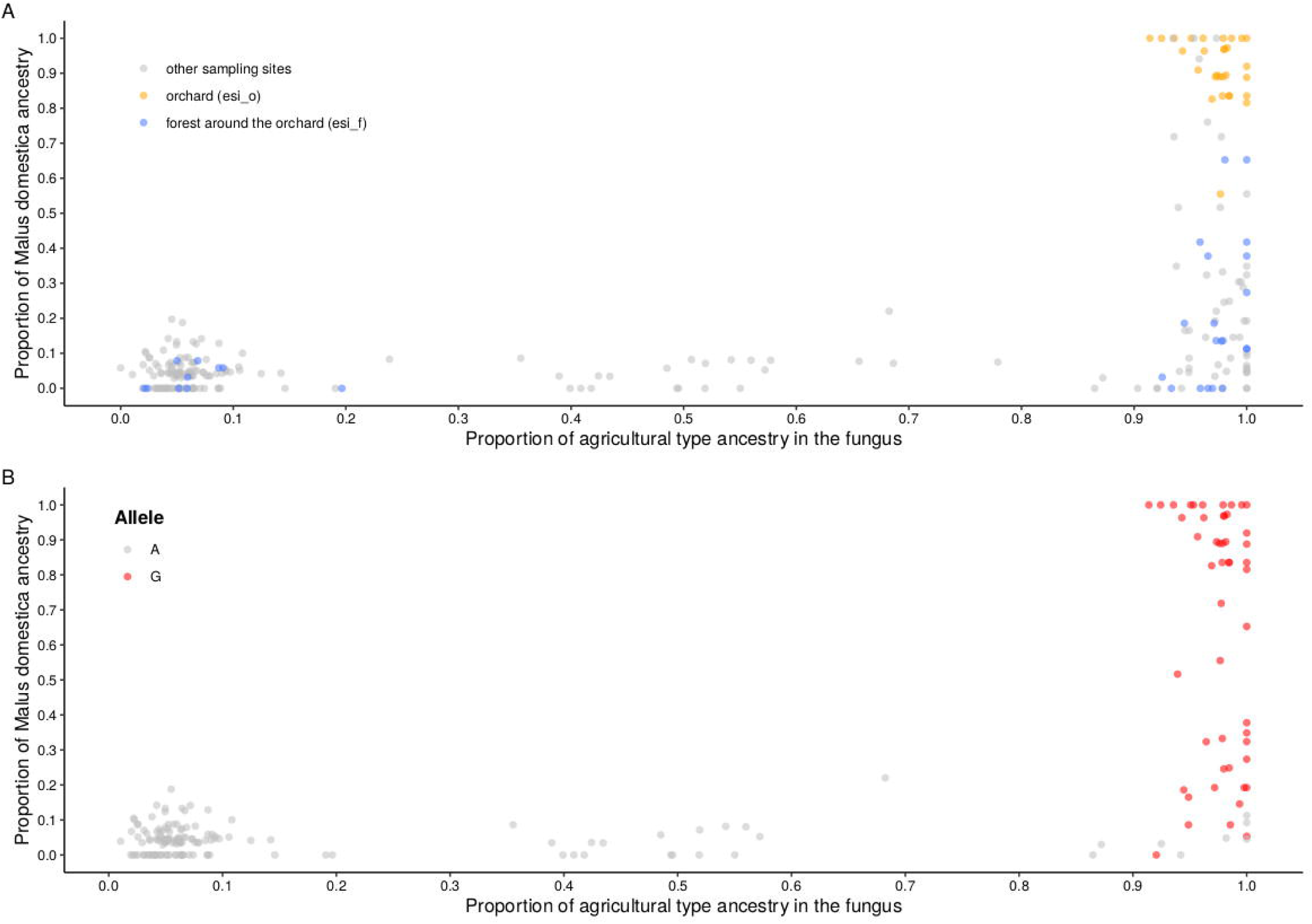
The proportion of *Malus domestica* ancestry in the apple tree (*y* axis) plotted against the proportion of agricultural-type ancestry in *Venturia inaequalis* strains (x axis). A) For each of the 249 host-pathogen pairs, two sampling sites are highlighted: the orchard (esi_o) in orange and the surrounding forest (esi_f) in blue. B) For each of the 189 host-pathogen pairs for which the genotype at the *V_081690_319* locus was available, the *V. inaequalis* strains carrying the tyrosine (A) allele were colored in grey and the ones carrying the stop codon (G) allele in red.

### Genetic determinants of host adaptation in *Venturia inaequalis*

In order to identify SNPs predicting the ability of fungal genotypes to parasitize the host species, we performed a DAPC on the genotypes of the 249 *V. inaequalis* strains, with groups defined *a priori* based on the assignment of their collection tree as *M. domestica*, hybrid or *M. sieversii*. The first linear discriminant function (LD1) clearly separated the strains sampled on domesticated apple trees from those sampled on wild and hybrid apple trees (Figure S8). The second linear function separated strains found on hybrid apple trees from the ones sampled on *M. sieversii* (LD2, Figure S8). The analysis of allele contribution (loadings) to the first discriminant function revealed that the SNP *V_08160_319* was strongly involved in the genetic differentiation between strains parasitizing *M. domestica* and those found on hybrid and wild hosts (loading of 0.123, Figure S9), while the other SNPs showed much lower contributions (from 8.723×10^−8^ to 0.024). The estimate of global *F*_*ST*_ was accordingly much higher for the V_08160_319 locus (*F*_*ST*_ = 0.887) than for other loci (Figure S10).

The SNP *V_08160_319* was located in a gene identified as a small secreted protein (SSP) using the protocol described in Le Cam et al. (2019). The two alleles of this SNP were A and G and corresponded to a non-synonymous site. According to the gene allelic sequences in the 33 reference genomes, the *V_08160_319* SNP was associated to a second one in the same codon, which thus corresponded to a TGA stop codon versus a TAC codon, coding for a tyrosine, at the 140th codon in the 295 amino-acid long protein. Out of the 189 strains for which genotypes for this SNP were available, the stop codon allele indeed had frequencies of 1.00 on *M. domestica* (N=28), 0.92 on hybrid apple trees (N=13) and 0.06 on *M. sieversii* (N=148) (Figure 4B). Considering ancestry of fungal strains, the stop codon allele had a frequency of 0.89 (N=56) in the agricultural-type population and 0 in the wild-type population (N=107) and in hybrids (N=26). The detection of only A (tyrosine) alleles in hybrids significantly deviated from expectations under the hypothesis of neutral segregation at this locus given the allele frequencies in the pure populations (Binomial-test, p-value=6.10×10^−5^).

To assess which allele was ancestral, we used five available genomes (Le Cam et al., 2019) belonging to the outgroup *V. inaequalis* lineage parasitising Pyracantha (Gladieux et al., 2010a; Le Cam et al., 2002). These genomes carried the TAC codon, as did the wild-type *V. inaequalis* population on *M. sieversii*. This means that the stop codon truncating the small secreted protein is a derived allele, and has evolved in the agricultural *V. inaequalis* population on *M. domestica*.

### Agricultural-type and hybrid fungal strains are more virulent than wild-type strains

We then investigated experimentally whether the observed association between fungal pathogen types and host tree species resulted from differences in infection ability, by inoculating 57 fungal strains (20 agricultural-type, 17 wild-type and 20 hybrid strains; Table S1) on one genotype of *M. sieversii* and one genotype of *M. domestica*. All agricultural-type strains could cause disease on *M. domestica* (N=20; Figures 5A and 6A), whereas only one wild-type and two hybrid strains could cause disease on *M. domestica*, and even then, only with very low levels of virulence (less than 2% of diseased leaf area, Figure 5A). All agricultural strains (N=20), hybrids (N=20) and wild strains (N=17) could cause disease on *M. sieversii* (Figure 5B). Both agricultural-type and hybrid strains were significantly more virulent (i.e. caused larger and/or more lesions) than wild-type strains on *M. sieversii* (Figures 5B and 6B; Wilcoxon tests: *P*=1.7 × 10^−5^ and *P*=1.2 × 10^−4^, respectively). The *V. inaequalis* hybrids displayed virulence levels on *M. sieversii* that appeared intermediate between the agricultural-type and wild-type populations although the difference between hybrids and the agricultural-type population was not significant (Figure 5B). Similar results were obtained when assignment to species and hybrids were done using the same hybrid index threshold as for apple trees (wild type: *Pagr*<0.2; hybrids: 0.2<*Pagr*<0.8; agricultural: *Pagr*>0.8; Figure S11). We found no significant correlation between the proportion of agricultural-type ancestry and the degree of virulence in the hybrid fungal strains (Figure S12; Kendall test: τ=-0.047, *P*=0.7702).

**Figure 5:**
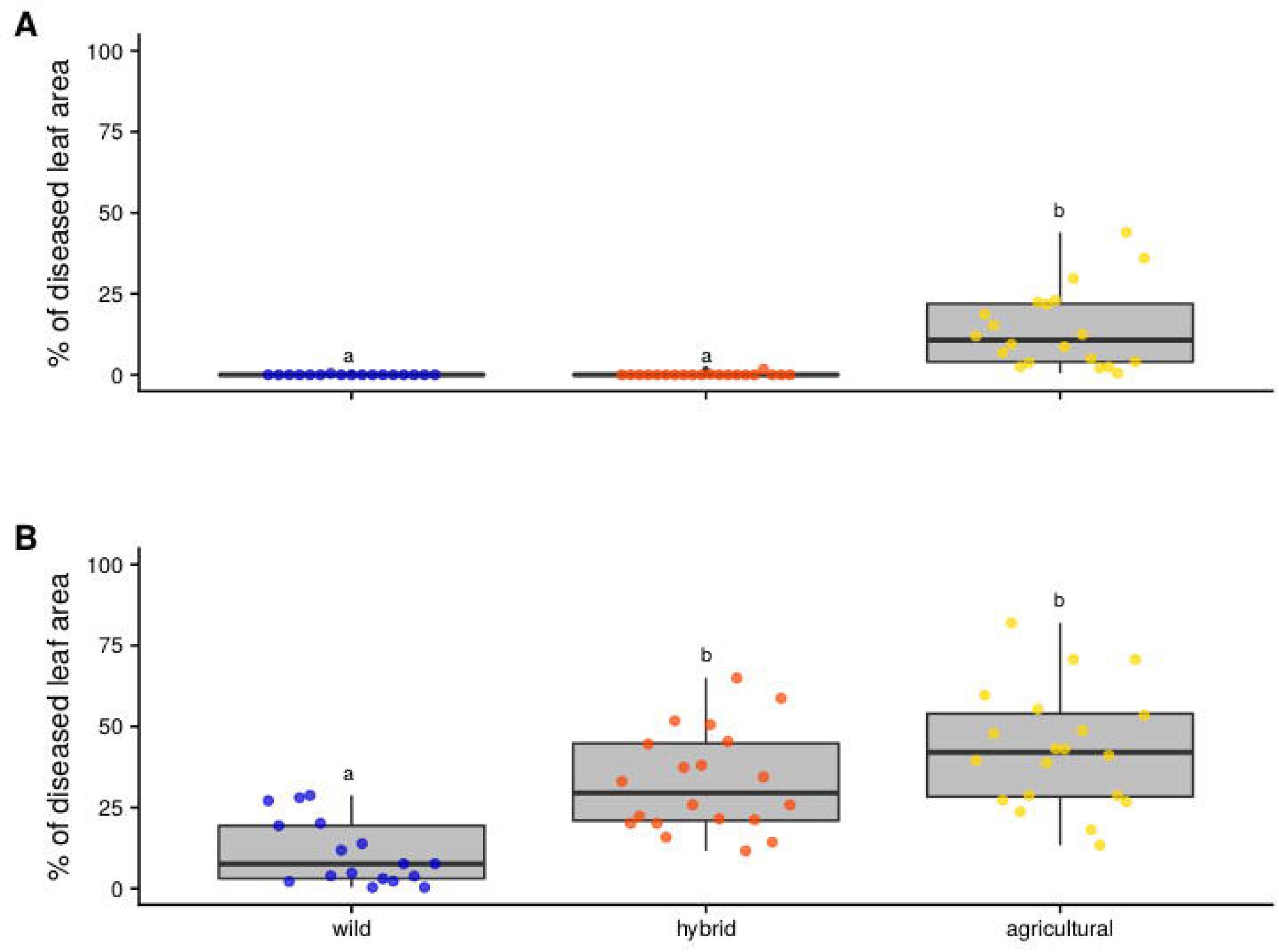
Results of pathogenicity experiments in controlled conditions consisting in artificial inoculation of the fungus *Venturia inaequalis* (57 fungal strains; mean of three replicates per strain, one genotype per apple species) on apple tree leaves (*Malus sieversii* and *M. domestica*). Boxplots of the percentage of scabbed leaf area for wild-type, agricultural-type and hybrid *V. inaequalis* strains of A) *M. domestica* at 21 days post inoculation (dpi) and B) *M. sieversii* at 19 dpi. The box represents the lower and the upper quartiles. The thick horizontal line represents the median. The whiskers represent the largest and lowest observed values that fall within the distance of 1.5 times the interquartile range. The points represent the mean values of the percentage of diseased leaf area for each strain across the three replicates. Different letters indicate significant differences between populations (*P*<0.05; Wilcoxon’s rank sum tests).

**Figure 6:**
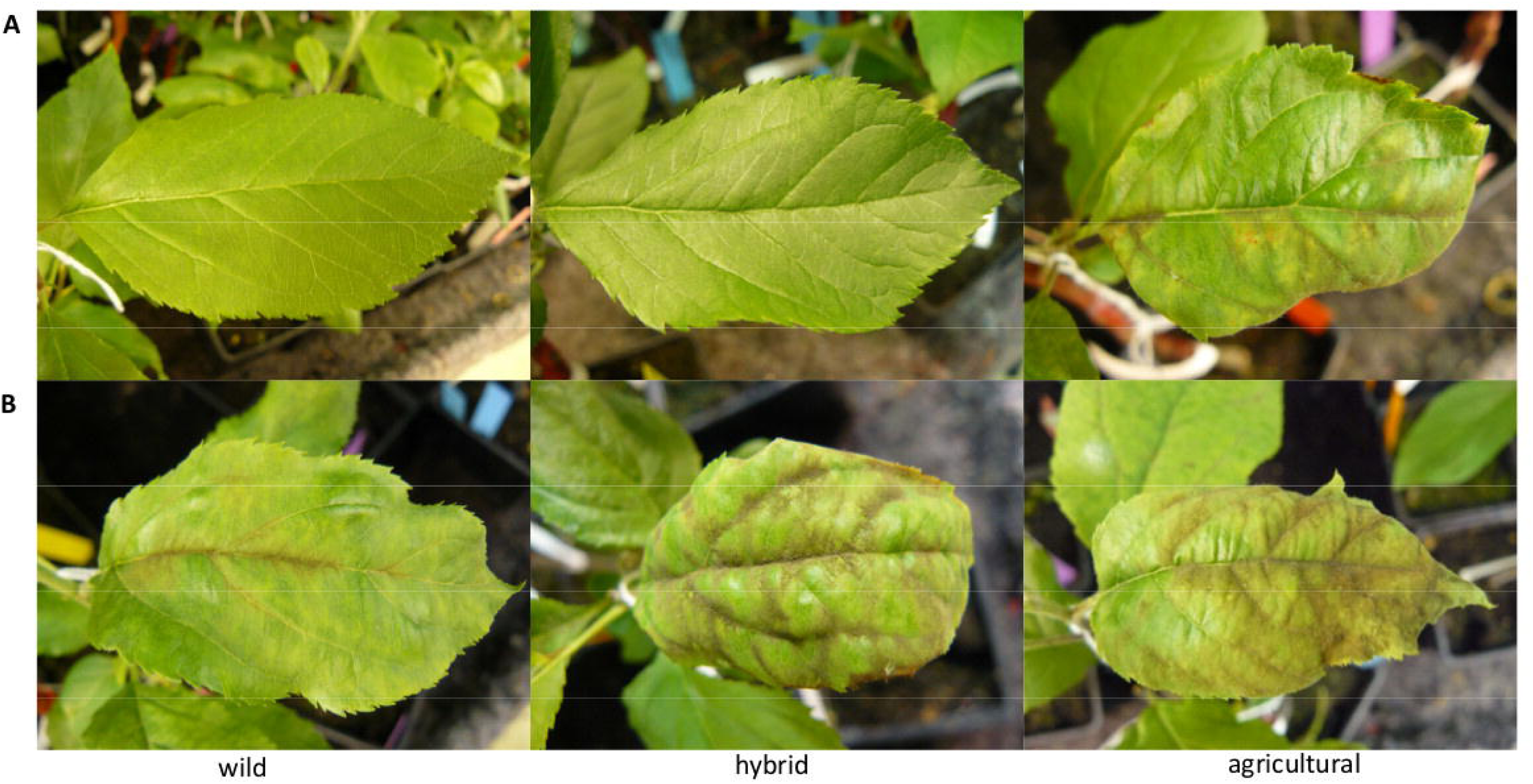
Pictures illustrating the percentage of scabbed leaf area for the most virulent wild-type, agricultural-type and hybrid *Venturia inaequalis* strains on A) *Malus domestica* at 21 days post inoculation (dpi) and B) *M. sieversii* at 19 dpi.

The allele at the outlier locus *V_081690_319* partially explained the virulence on *M. domestica*. Fourty-seven genotypes were available out of 57 strains used in the experiment. Out of the 15 agricultural-type strains with available genotypes, 13 (87%) carried the stop codon (TGA) allele. Conversely, all the 15 hybrid strains with available genotypes (100%), as well as all the 17 wild strains (100%), carried the tyrosine (TAC) allele. When comparing the two groups of strains corresponding to their alleles at the *V_081690_319* locus, we found that the strains carrying the stop codon (TGA) allele were more virulent than those carrying the tyrosine (TGA) allele, both on *M. domestica* and on *M. sieversii* (Wilcoxon tests: *P*=1.4×10^−8^ and *P*=1.2×10^−3^, respectively; Figure S13). Most of the strains carrying the tyrosine allele could not cause disease at all on *M. domestica*, but a few of them could induce a low level of symptoms (Figure S13A).

### Demographic inference on the divergence history of wild-type and agricultural-type *Venturia inaequalis* populations

We performed demographic inferences in order to test if the observed co-occurrence and hybridization of wild and agricultural-type *V. inaequalis* populations in Kazakhstan resulted from divergence with gene flow or originated from a secondary contact, following the introduction of *M. domestica*, about one century ago in a few orchards in forests of wild *M. sieversii*. We compared the likelihoods of four contrasted scenarios of divergence, with either strict isolation, isolation with continuous migration, isolation with initial migration or secondary contact (Figure 7A).

**Figure 7:**
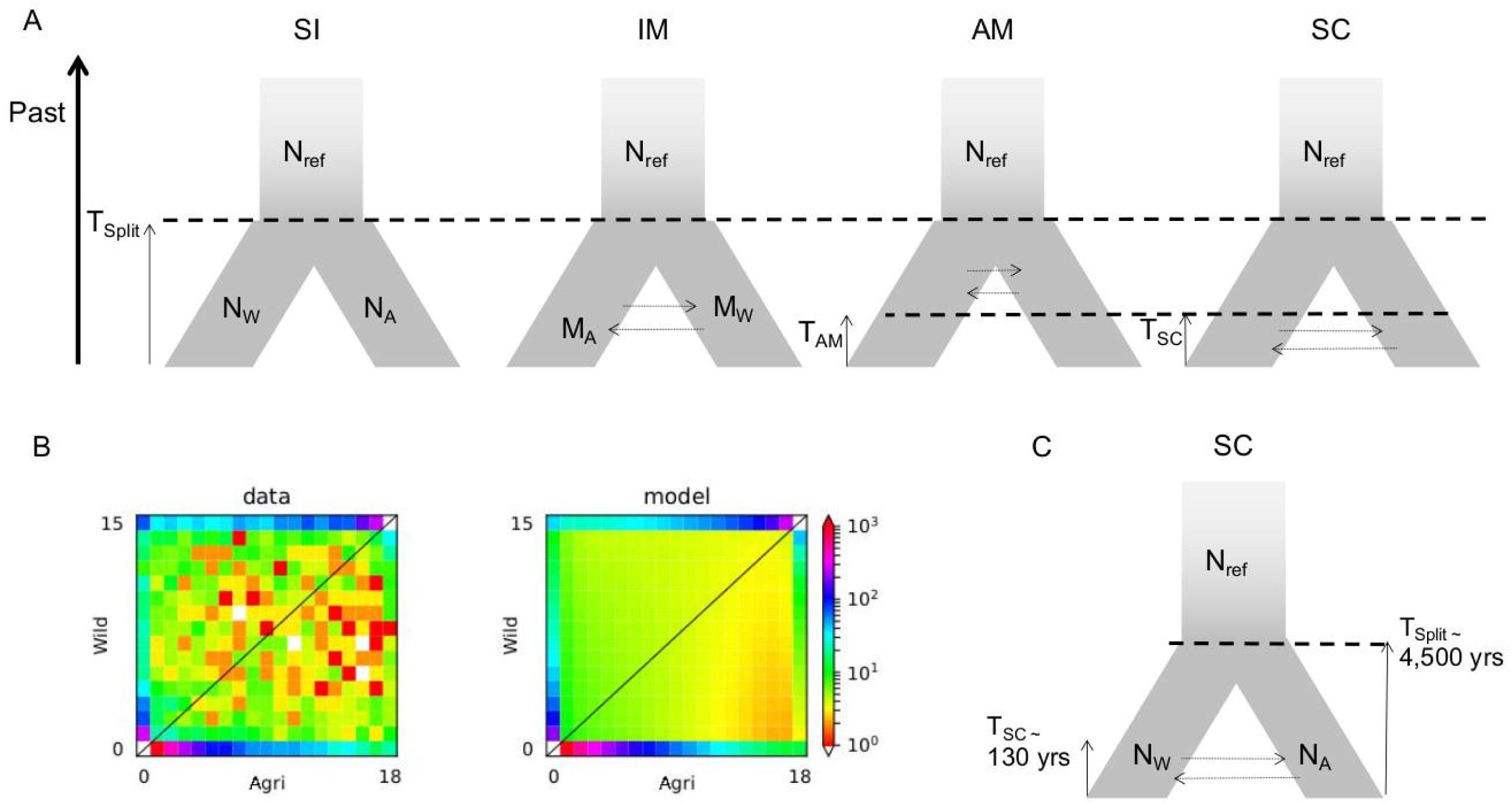
Demographic inference on the divergence between wild and agricultural types of *Venturia inaequalis* in Kazakhstan. **A**. The four contrasted scenarios of divergence compared, with either strict isolation (SI), isolation with continuous migration (IM), isolation with initial migration (AM) and secondary contact (SC), with effective populations sizes of the ancestral population (N_ref_) and of the two daughter populations (N_w_ and N_A_), the rate of gene flow in the two directions (M_w_ and M_A_) and the time since the cessation of gene flow (T_AM_) or the secondary contact (T_SC_). **B**. The joint allele-frequency spectrum (AFS) for the wild and agricultural fungal populations, showing the count of derived allele. Each entry of the joint AFS is colored by the number of SNPs in it, according to the scale shown. **C**. The secondary contact (SC) model that obtained the best support and estimates of the parameters.

The models allowing gene flow received better support than the strict isolation model (AIC= 2403,587014; LogL= −1197,793507). The model of isolation with migration (IM; AIC= 1975,638252; LogL= −981,8191258) did not perform significantly better than the ancient migration model (AM; AIC= 1977,698045; LogL= −981,8490226). The secondary contact model (SC) received the best support (AIC= 1871,313317, LogL= −928,6566583), which indicated that the agricultural-type *V. inaequalis* population was likely absent in Asian wild apple forests before the recent introduction of domesticated apple trees in orchards (Figure 7B; Table 1). Using the estimate of θ obtained from the best model (SC, θ=143.729), a mutation rate μ of 2×10^−8^, we found that the ancestral effective size before split, *N*_*ref*_, was near 20,000 individuals (Figure 7C). Parameter estimates for the secondary contact model indicated that the divergence between the wild-type and agricultural-type *V. inaequalis* populations occurred 4,570 years ago (4,570 ± 324 SD) followed by a secondary contact 130 years ago (130 ± 9 SD). This scenario and estimates are consistent with a fungal divergence triggered by apple domestication under strict genetic isolation, followed by a recent secondary contact promoted by modern orchards (Figure 7C).

**Table 1:** Comparison of four alternative demographic models of divergence between the reference wild-type and agricultural-type populations of *Venturia inaequalis* using ∂a∂i on the SNP dataset.

| Model | $k$ | logL | AIC | Model Comparison | $N_{\text{Wild}}$ | $N_{\text{Agri}}$ | $M_{\text{WA}}$ | $M_{\text{AW}}$ | $T_{\text{S}}$ | $T_{\text{AM}}$ | $T_{\text{SC}}$ |
| --- | --- | --- | --- | --- | --- | --- | --- | --- | --- | --- | --- |
| SI | 3 | -1197.8 | 2403.587014 | - | 0.094 | 1.360 | - | - | 0.092 | - | - |
| IM | 5 | -981.8 | 1975.638252 | SI*** | 0.139 | 0.898 | 1.259 | 1.131 | 0.460 | - | - |
| AM | 6 | -981.8 | 1977.698045 | SI***, IM<sup>NS</sup> | 0.139 | 0.892 | 1.270 | 1.140 | 0.460 | 1.09x10<sup>-9</sup> | - |
| **SC** | **6** | **-928.6** | **1871.313317** | **SI\*\*\*, IM\*\*\*, AM<sup>+++</sup>** | **0.163** | **0.991** | **1.355** | **4.389** | **0.227** | - | **0.006** |

SI, strict isolation model; IM, isolation with migration model; AM, ancient migration model; SC, secondary contact model; $k$ , parameter numbers; logL, log-likelihood; AIC, Akaike Information Criterion; $N_{\text{Wild}}$ , effective sizes of wild-type lineage; $N_{\text{Agri}}$ , effective sizes of agricultural-type lineage; $M_{\text{WA}}$ , migration rates from wild to agricultural-type lineage; $M_{\text{AW}}$ , migration rates from agricultural-type to wild-type lineage; $T_{\text{S}}$ , time of split; $T_{\text{AM}}$ , duration time since ancient migration; $T_{\text{SC}}$ , duration time since secondary contact.
The best supported model (SC) is indicated in bold. Model comparisons within and between classes of models are shown. Nested models were compared using likelihood ratio tests, with subscripts indicating significance levels (abbreviated \*\*\* $P < 0.001$ ; \*\* $P < 0.01$ ; \* $P < 0.05$ , NS non-significant). Non-nested models were compared using AIC with relative likelihood of each model compared to the best model $L(\text{Mi}|\text{Mbest}) = \exp((\text{AICmin} - \text{AICi})/2)$ (abbreviated +++ $L(\text{Mi}|\text{Mbest}) < 0.001$ ; ++ $L(\text{Mi}|\text{Mbest}) < 0.01$ ; + $L(\text{Mi}|\text{Mbest}) < 0.05$ , $L(\text{Mi}|\text{Mbest}) > 0.05$ : AIC difference shown).

## Discussion

### Gene flow between apple trees species may threaten the genetic integrity of the Asian wild apple

Our data show that *M. sieversii* wild apple trees are affected by gene flow following the introduction of the domesticated apple trees in orchards near the natural apple forests of the Kazakh Tian Shan Mountains. Gene flow from *M. domestica* to *M. sieversii* had previously been suggested (Cornille et al. 2012; Omasheva et al., 2017), and we reveal here the strong impact of the presence of orchards close to wild forests. Currently, gene flow appears to pose a less severe threat to *M. sieversii* than it does to *M. sylvestris* in Europe, where wild European crabapple tree populations display massive introgression from the cultivated apple tree, to the extent that the European wild species is considered as endangered (Cornille et al., 2015; Feurtey et al., 2017). The lower level of introgression in *M. sieversii* wild populations in Central Asia is probably due to more recent and much less extensive secondary contact between wild and cultivated species than in Europe. The cultivated apple tree was introduced into Europe before the 3^rd^ century BC (Cornille et al., 2014), whereas it was not introduced into the area sampled in this study until about 100 years ago, and much less extensively than in Europe. Furthermore, the higher density of *M. sieversii* trees in Kazakh wild forests (Harris et al., 2002; Vavilov, 1931) than of *M. sylvestris* in European wild forests probably also restricts gene flow from cultivated apple trees. Indeed, pollination distances decrease with density around pollinated trees in European wild apple trees, longer pollination distances increasing the risk of interspecific mating with cultivated trees (Feurtey et al. 2017; Reim et al. 2015).

While the overall percentage of apple tree hybrids in the wild forest remained low, near 25% of the trees were found to be hybrids in the forest immediately adjacent to a cultivated orchard, as well as in the forest location that was the closest to the peri-urban environment of Almaty. These findings confirm the weak interspecific reproductive barriers reported for the *Malus* genus (Zohary & Hopf, 2000) and raise concerns about the consequences of increasing human-induced changes in this region. As indicated by the IUCN Red List, *M. sieversii* is already threatened by its range shrinking in Central Asia (Eastwood, Lazkov & Newton, 2009), with a disappearance of more than 70% of the apple wild forest area over the last 30 years due to agricultural expansion and overgrazing. In addition to this range reduction, our results support the view that the genetic integrity of *M. sieversii* may be also threatened. Thus, unless preventive measures are taken to protect Central Asian forests, it may only be a question of time before the genetic integrity of *M. sieversii* deteriorates in Central Asia, as already observed for *M. sylvestris* in Europe (Cornille et al., 2015; Feurtey et al., 2017), as well as for other European native plant species such as poplars (Bleeker, Schmitz & Ristow, 2007; Rhymer & Simberloff, 1996; Vanden Broeck, Villar, Van Bockstaele & Van Slycken, 2005).

### Recent invasion of the wild forests by the agricultural *V. inaequalis* population

We confirm in this study the existence of two differentiated *V. inaequalis* populations in Central Asia. The high differentiation level between the agricultural-type and wild-type *V. inaequalis* populations raises the question of whether they should be considered as separate species. However, further studies are needed to assess the taxonomic status of the *V. inaequalis* lineages, especially as hybrids were found.

Most importantly, we show that agricultural-type and wild-type *V. inaequalis* populations came into a recent secondary contact after having diverged in strict isolation during apple tree domestication. Indeed, demographic inferences showed that agricultural-type and wild-type fungal populations initially diverged without gene flow about 4,500 years ago, a time consistent with that of apple tree domestication, and came into secondary contact about one century ago, i.e., when domesticated apple trees were introduced in Central Asia. This secondary contact has led to the recent invasion of the agricultural-type population in the wild *M. sieversii* forest and to the production of hybrids.

### The agricultural-type *V. inaequalis* population is able to parasitize both wild and domesticated host trees

We uncovered contrasting host range patterns among *V. inaequalis* populations: wild-type and hybrid fungal strains were only found on *M. sieversii* trees, whereas agricultural-type fungal strains were found on trees of all types of ancestry. Pathogenicity tests in controlled conditions matched the distribution observed in nature. Indeed, the agricultural-type fungal strains were able to cause disease on *M. domestica* and *M. sieversii* whereas the wild-type and hybrid fungal strains could induce symptoms only on wild host trees. These results, obtained with high spore concentration and in favorable climatic conditions, are conservative and confirm conclusions that were previously obtained with a limited number of *V. inaequalis* strains (Lê Van et al., 2012). Furthermore, we detected agricultural-type *V. inaequalis* strains at eight different sites in the wild forests on numerous different trees, suggesting that the agricultural-type population can parasitize a large range of *M. sieversii* genotypes. In contrast, no wild-type or hybrid fungal strain was detected on any of the *M. domestica* genotypes present in the orchard. This host range pattern likely explains the significant correlation found between apple tree and *V. inaequalis* pairwise *F*_*ST*_ matrices, which was not due to isolation by distance patterns in hosts and pathogens. Indeed, it is likely that the *M. domestica* ancestry proportion in each site drives the allele frequency distribution in *V. inaequalis* by selection, because *M. domestica* can only be parasitized by agricultural-type strains, and not by wild-type or hybrid pathogens. This situation contrasts with that reported for poplars, in which the hybrid fungal pathogen *Melampsora x columbiana* can parasitize both pure species and hybrid trees (Newcombe et al., 2000).

In addition, the agricultural *V. inaequalis* population and hybrids displayed greater virulence on wild apple trees than its own endemic wild-type population. This is consistent with the process known as “pestification” (Saleh, Milazzo, Adreit, Fournier & Tharreau, 2014), under which selection by humans of more resistant plants unwittingly leads pathogens to accumulate more virulences traits and to cause more severe symptoms. This could add further threats for the wild tree species in its natural habitat. Indeed, the scab disease on apple tree leaves reduces photosynthesis (Spotts & Ferree, 1979), which may affect the growth of the plants. While such increase in virulence may have a limited impact on the growth of adult trees, its effect on the growth of young seedlings could be severe. Furthermore, scab disease can lead to early fruit fall (MacHardy, 1996), which may decrease seed number, thereby affecting host population dynamics. On *M. sieversii*, hybrid strains were more virulent than wild-type fungal strains but not significantly different from agricultural-type fungal strains. Thus, although the hybridization between fungal lineages does not currently appear to increase damages on wild trees compared to the pure agricultural pathogen, it might participate in the invasion of the disease on the wild host. Future experiments are still needed to estimate precisely the disease fitness effect on seedlings and on the number of seeds produced by adult trees in order to ascertain the long-term impact of this pathogen population on the wild tree natural populations. Nevertheless, our results already suggest that the invasion of the wild forests by the agricultural-type *V. inaequalis* population could further threaten natural *M. sieversii* populations.

### The genomic determinants of pestification: a likely new virulence gene

Our findings suggest the evolution of *V. inaequalis* populations by host tracking of the domesticated apple tree without the loss of its ability to parasitize the wild host. Many resistance genes have been introgressed into *M. domestica* during domestication and modern breeding (Cornille et al., 2012). During its evolutionary tracking of apple trees evolution, the agricultural *V. inaequalis* population has probably accumulated multiple alleles allowing to counteract the crop resistance genes, potentially including the truncated allele of the small secreted protein identified in this study as a putative avirulence gene. Indeed, we found that the host lineage on which *V. inaequalis* strains were sampled was predicted by a SNP in a gene encoding a small secreted protein. In the fungal strains collected on *M. domestica*, the allele corresponded to a stop codon while strains sampled on *M. sieversii* carried the full-length gene. Taken together with the presence of the full-length gene in an outgroup, the *V. inaequalis* lineage parasitizing Pyracantha (Gladieux et al., 2010a; Le Cam et al., 2002), this suggests the acquisition of a virulence allele by the agricultural *V. inaequalis* population during crop host tracking, through the loss of a protein recognized by the domesticated apple tree selected for resistance to apple scab. Such an increase in the virulence of a pathogen following domestication of its host is compatible with the pestification process (Salehet al., 2014), and is consistent with virulence evolution mechanisms previously found in many crop pathogens (Raffaele & Kamoun, 2012).

The identification of a putative avirulence gene from a small set of 181 SNPs can be explained by the initial choice of these SNPs as the most differentiated between the fungal populations and located within genes. However, we cannot exclude that the *V_081690_319* SNP is not directly involved in host adaptation but is instead linked to a genomic determinant of host adaptation. Future functional experiments are required to confirm that the identified gene is an avirulence gene. On the plant side, the identification of the receptor involved in the recognition of this putative avirulent protein could lead to the discovery of a new resistance gene.

More comprehensive studies on whole *V. inaequalis* genomes will likely detect further SNPs involved in host adaptation, as suggested by our data. Indeed, the SNP allele corresponding to the stop codon (G allele) was not found in any of the 15 hybrids for which genotypes were available, which significantly deviated from neutral expectations, while agricultural-type strains carrying this allele were found on *M. sieversii*, even if it was at low frequency. In addition, a few strains carrying the full-length allele of the putative avirulence gene were able to cause some symptoms on *M. domestica* in artificial inoculations. This suggests that the stop codon allele was not selected against on *M. sieversii* in itself but may be deleterious in association with alleles at other loci in hybrids. These results therefore suggest that the locus *V_081690_319* is involved in epistatic interactions with other genes and that hybridization breaks up beneficial allelic combinations required to parasitize *M. domestica*. Furthermore, the *V. inaequalis* hybrids appeared quantitatively intermediate in virulence on *M. sieversii*, although the virulence difference with the agricultural population was not significant. Intermediate values in hybrids for haploid organisms suggest the implication of multiple genes in virulence.

## Conclusion

We found that natural *M. sieversii* populations are currently only mildly affected by gene flow from *M. domestica*, which was introduced into the area about a century ago but still remains very rare in the Kazakh mountains. Further disturbances in this area might however lead to much higher levels of crop-to-wild gene flow, as already reported in Europe for the European crabapple (Cornille et al., 2013; 2015; Feurtey et al. 2017). In addition, the agricultural-type *V. inaequalis* population is invading wild forests and is introgressing the wild-type *V. inaequalis* population, probably due to the greater virulence of the agricultural-type fungal population acquired during its tracking of apple tree domestication. Furthermore, our study represents one of the very rare joint analyses of host and pathogen populations (Croll & Laine, 2016), despite their importance for understanding the evolutionary mechanisms and histories leading to host specialization and local adaptation. Thanks to the study of host and pathogen pairs, we could identify a putative avirulence gene that may be of great importance on a cultivated crop, and a similar approach may be widely used in future studies on host and pathogen populations.

## Acknowledgments

The authors thank C. Peix and kazakh forestry agents for their assistance in sampling, M. Cascales for her assistance in SNPs data analyzing, N. Bierne, R. Nielsen and M. Slatkin for helpful discussions. They acknowledge the staff at the GENTYANE genotyping platform (INRA GDEC, Clermont-Ferrand, France) for SNP and microsatellite genotyping. Authors are also grateful to Pascal Heitzler for providing *M. sieversii* budwoods. They are also most grateful to the PHENOTIC core facility (Angers, France) for its technical support. This study was funded by grants from ANR-12-ADAP-0009 (Gandalf project), from INRA-SPE Department (Escapades project), by a Louis D. grant (Institut de France) to TG, a PhD grant from the Région Ile-de-France to AF, and by the program “Objectif Végétal, Research, Education and Innovation in Pays de la Loire”, supported by the French Region Pays de la Loire, Angers Loire Métropole and the European Regional Development Fund (Escapes project).

## Data accessibility

All data will be archived and available on the following link, doi: 10.5281/zenodo.3565732

## Author Contributions

AF, EG, TG and CL wrote the manuscript with inputs from BLC, VC, JS and LD. Sampling in Kazakhstan was performed by CL and BLC. Apple microsatellite genotyping was performed by AF. Apple scab SNP design and genotyping was performed by MDG, MS and CL. SNP mapping and calling was performed by LD and JS. EG, MNB, PE and VC managed *in vitro* cultures of *Venturia inaequalis* and pathology assays. AF, EG and CL performed population genetics and statistical analyses. CL, BLC and TG conceived the project and obtained financial support.

## Notes

### Competing Interest Statement

The authors have declared no competing interest.

