## Supplementary material for "Asian wild apples threatened by gene flow from domesticated apples and by their pestified pathogen": Supp_Figures_and_tables

***Molecular Ecology* Supporting Information**

**Article acceptance date:** Click here to enter a date.

The following Supporting Information is available for this article:

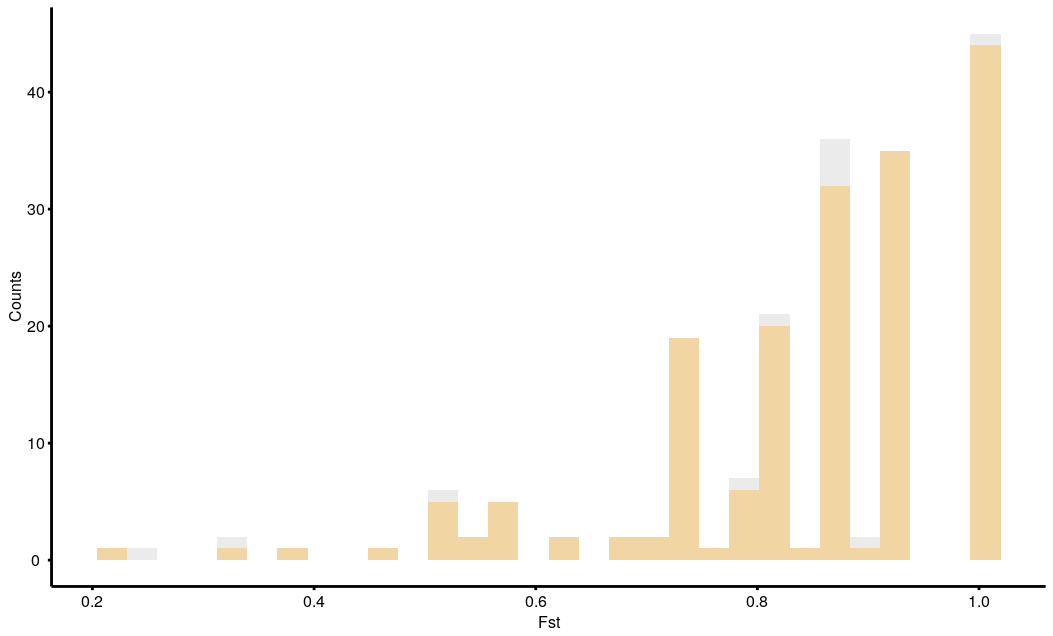

**Fig. S1:** Distribution of F_ST_ estimates between wild and agricultural-types *Venturia inaequalis* populations for all 192 SNPs (in grey) and the set of 181 SNPs finally retained (in orange).

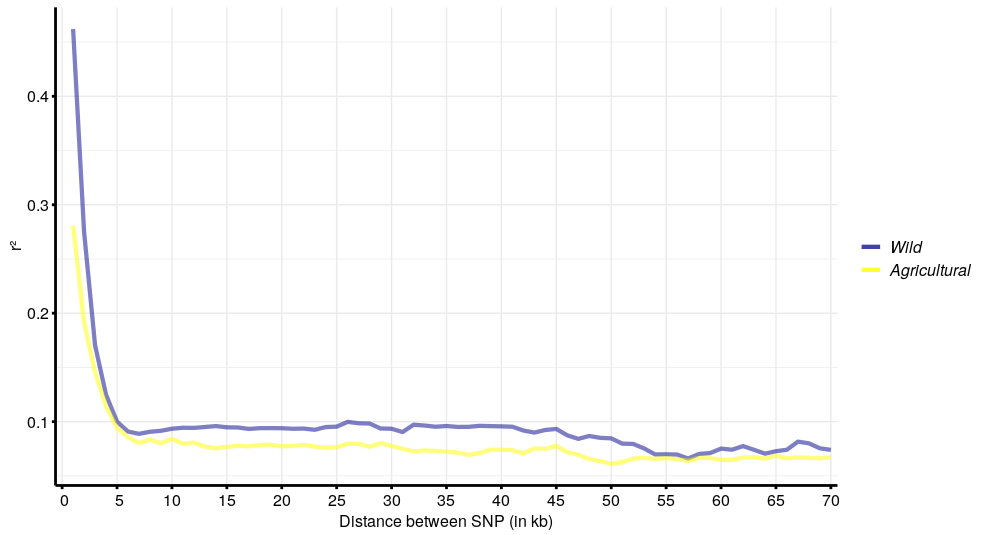

**Fig. S2:** Linkage disequilibrium (LD) decay as a relationship between the correlation coefficient between markers *(r²)* and the physical distance between markers in kilobases in *Venturia inaequalis* populations. LD decay is represented in blue for the wild population and in yellow for the agricultural one.

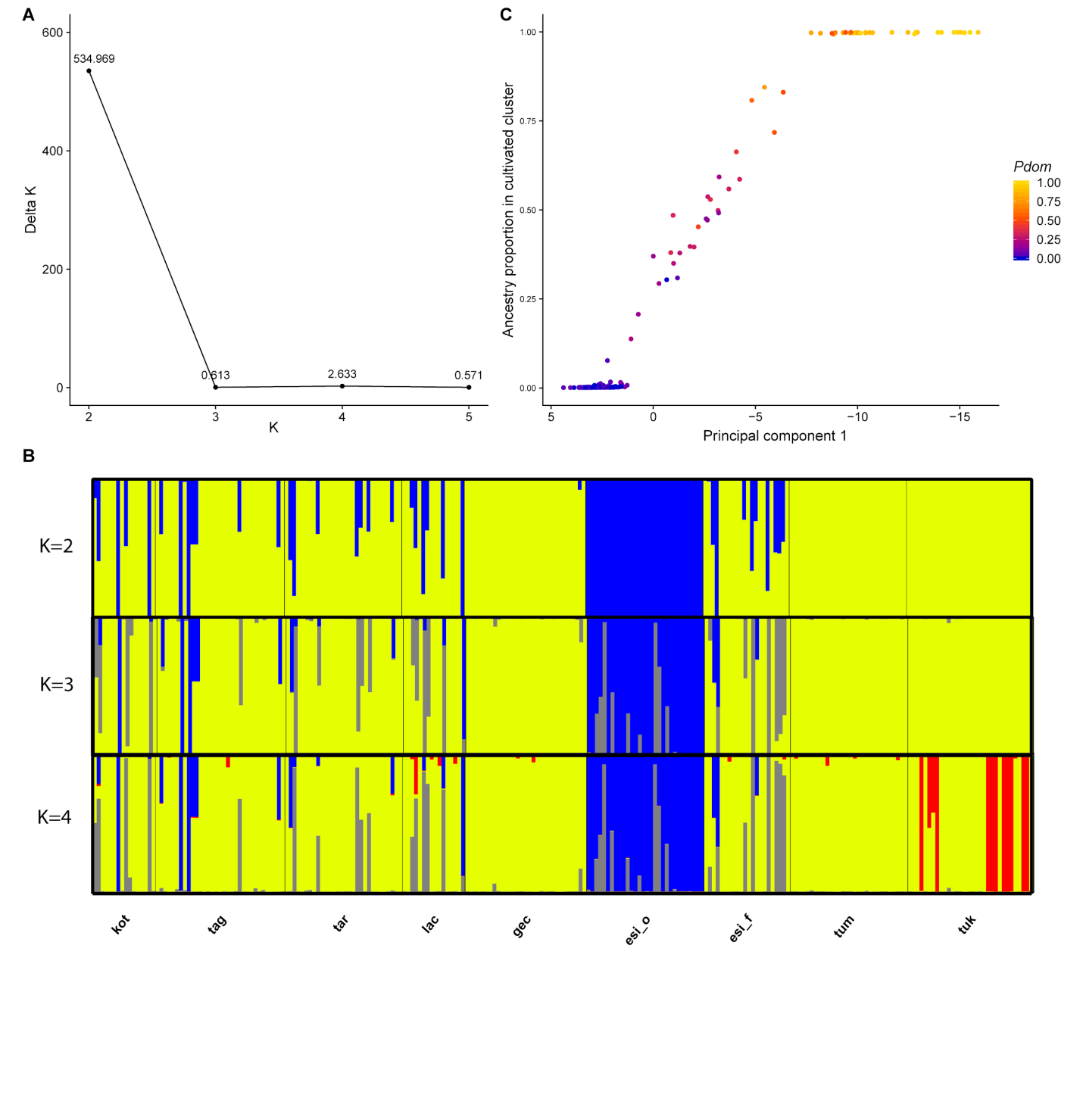

**Fig. S3:** Apple tree ancestry inferred from the Bayesian clustering implemented in STRUCTURE. A) Inference of the number of clusters corresponding to the strongest subdivision level (Evanno et al., 2005), corresponding to the highest value of deltaK. B) Proportion of ancestry of apple tree genotypes for *K* = 2 to 4 clusters inferred with the STRUCTURE software. Each vertical bar represents an individual and shows its assignment to the clusters. Individuals are grouped according to their sampling site. C) Relationship between the coordinates on the first axis of the principal component analysis (PCA) and the proportion of *Malus domestica* ancestry as inferred from STRUCTURE. Dots are colored according to the proportion of *M. domestica* ancestry *P_dom_.* For clarity, the PCA axis was reversed.

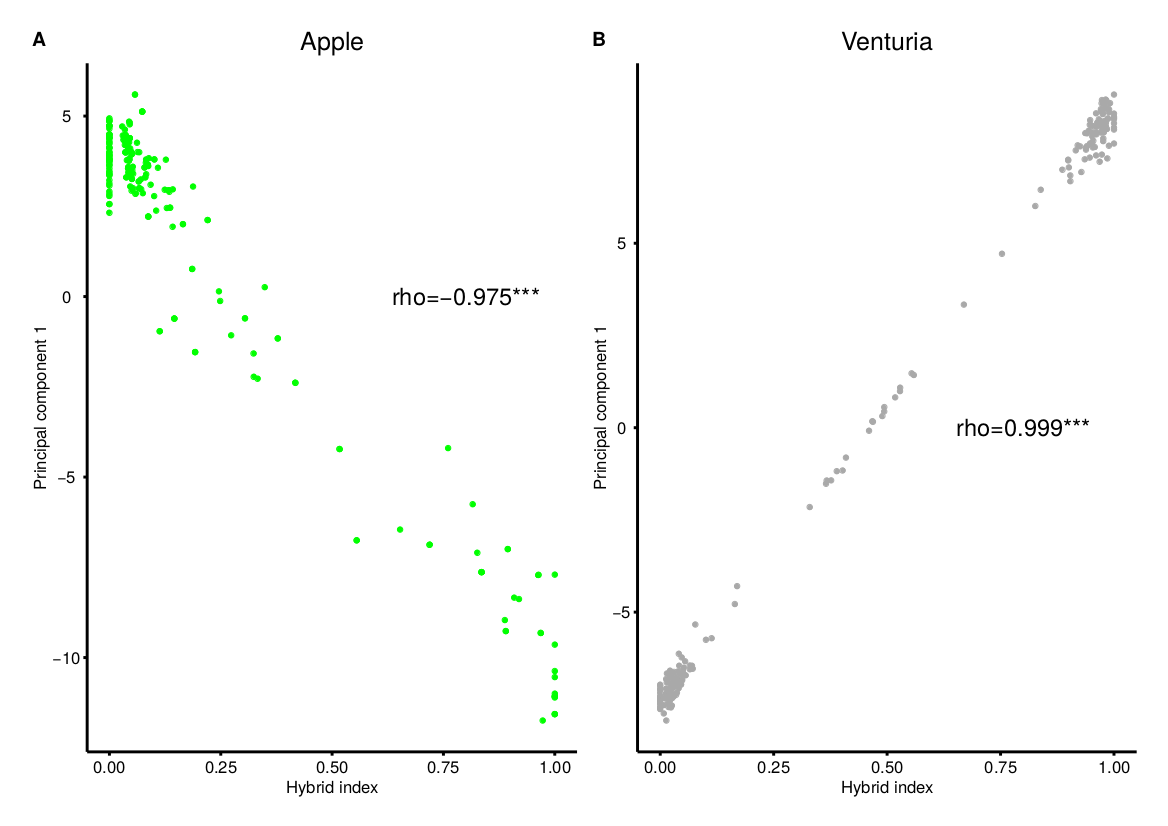

**Fig. S4:** Correlations between the hybrid index inferred from INTROGRESS and the coordinates on the first axis of the principal component analysis (PCA) for the apple trees (A) and *Venturia inaequalis* (B).

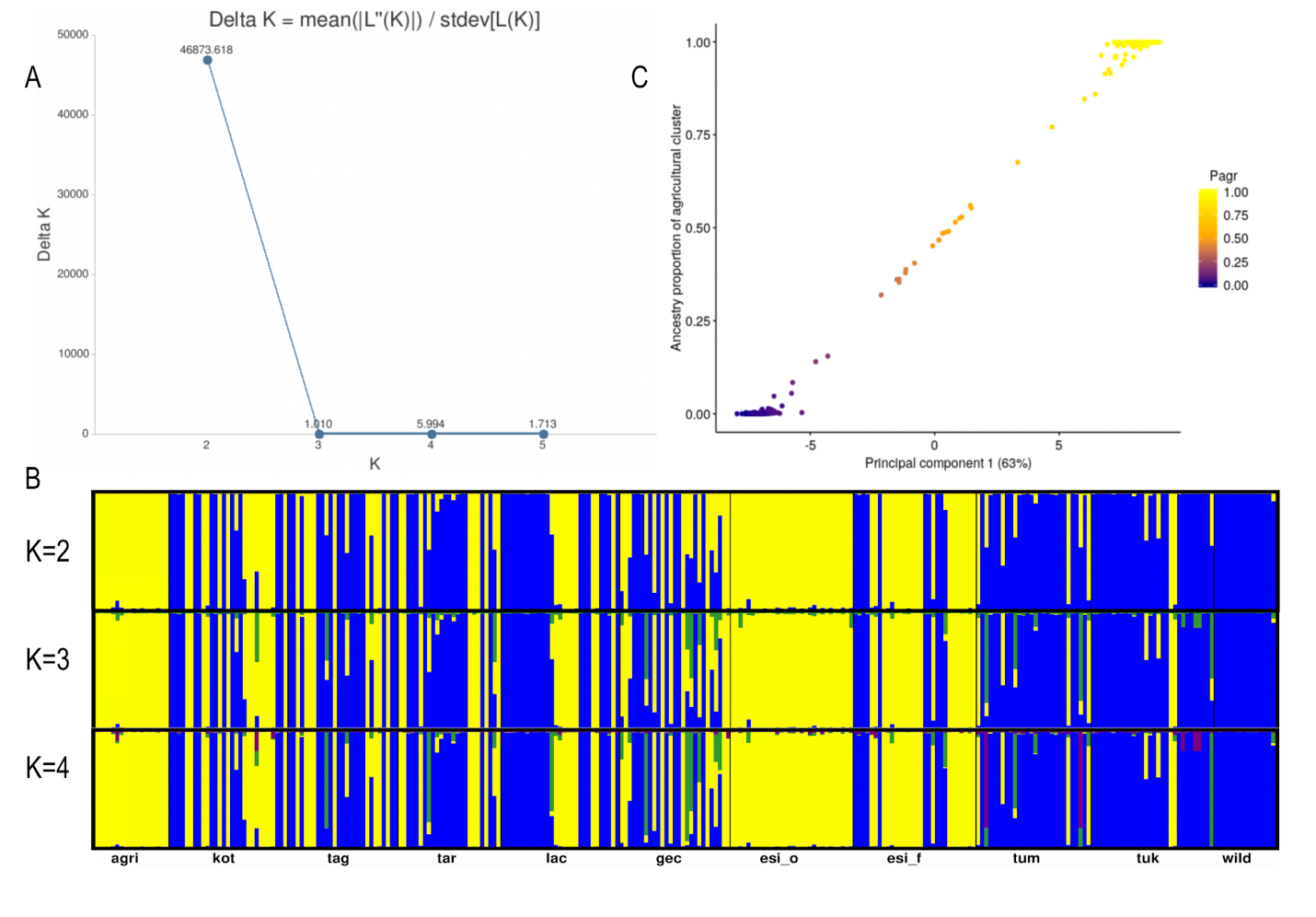

**Fig. S5:** *Venturia inaequalis* ancestry inferred from the Bayesian clustering implemented in STRUCTURE. A) Inference of the number of clusters corresponding to the strongest subdivision level using the method described in Evanno et al. (2005), corresponding to the highest value of deltaK, here K=2. B) Proportion of ancestry of apple tree genotypes for *K* = 2 to 4 clusters inferred with the STRUCTURE software. Each vertical bar represents an individual and shows its assignment to the two clusters. Individuals are grouped according to their sampling site. “Agri” and “wild” represent the reference populations. C) Relationship between the coordinates of the first axis of the principal component analysis (PCA) and the proportion of agricultural population ancestry as inferred from STRUCTURE. Dots are colored according to the proportion of agricultural ancestry *P_agr_.*

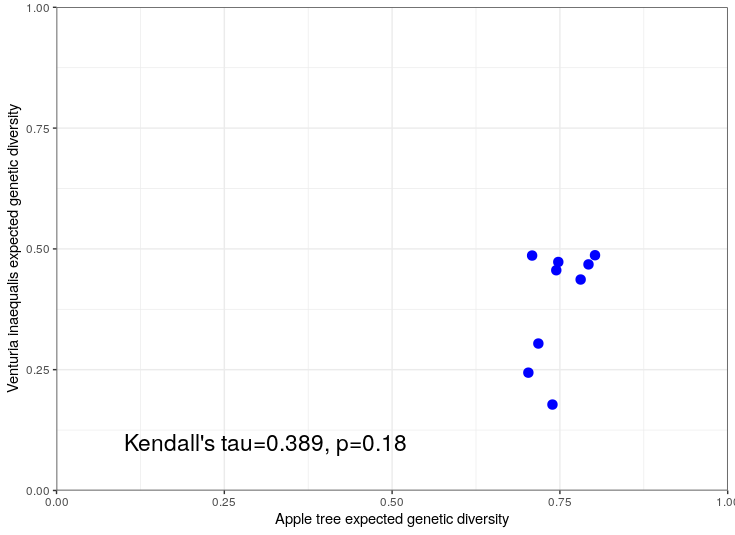

Fig. S6: Correlation analysis between genetic diversities in apple trees and in *Venturia inaequalis.* The result of Kendall’s tau correlation test shows no significant correlation.

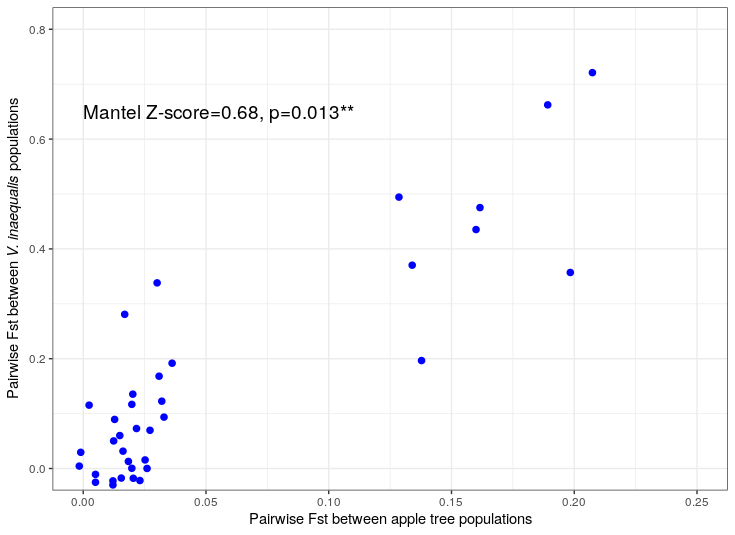

**Fig. S7:** Correlation between apple tree and *Venturia inaequalis* pairwise *F_ST_* matrices. The correlation is significant as indicated by the Mantel Z-score.

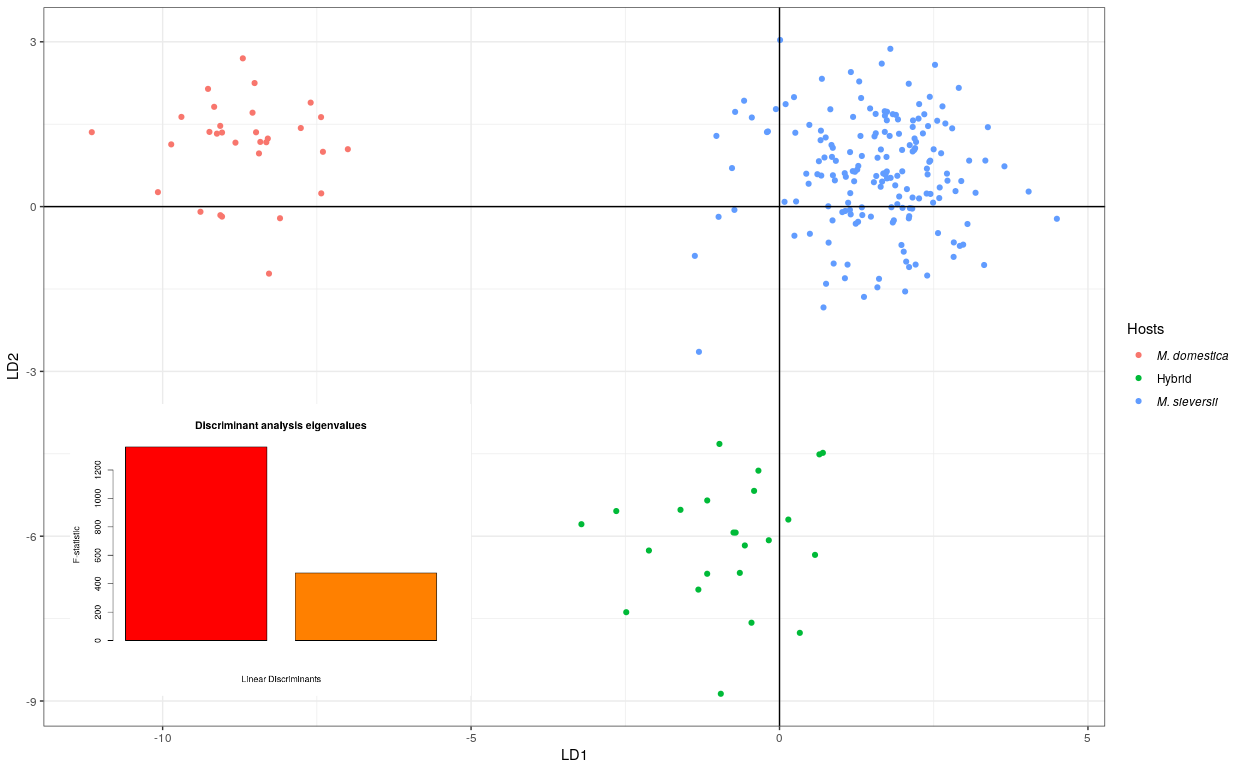

**Fig. S8:** Discriminant analysis of principal component (DAPC) of *Venturia inaequalis* specifying as *a priori* groups the assignment of their apple tree of collection to the apple populations (*Malus domestica* in red, *M. sieversii* in blue and hybrids in green). On the bottom left corner, the two discriminant functions eigenvalues are shown.

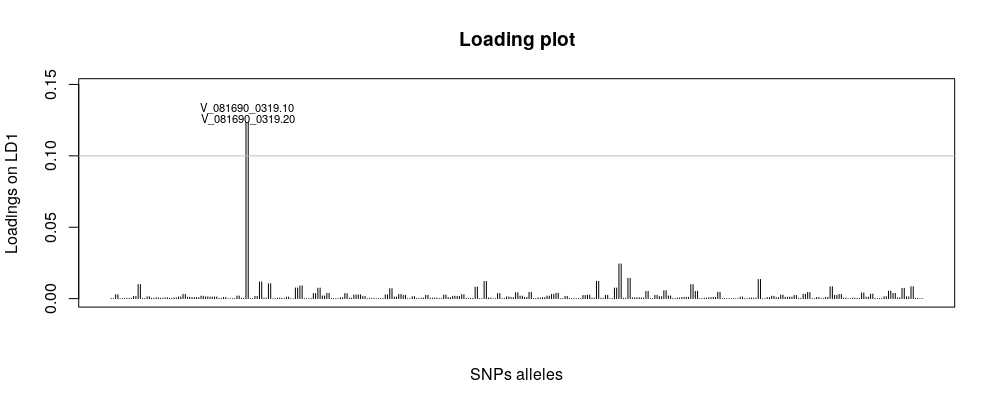

**Fig. S9:** Loadings on the first discriminant function of the discriminant analysis of principal component (DAPC). Each vertical bar represents a SNP. The horizontal line represents an arbitrary threshold for significance.

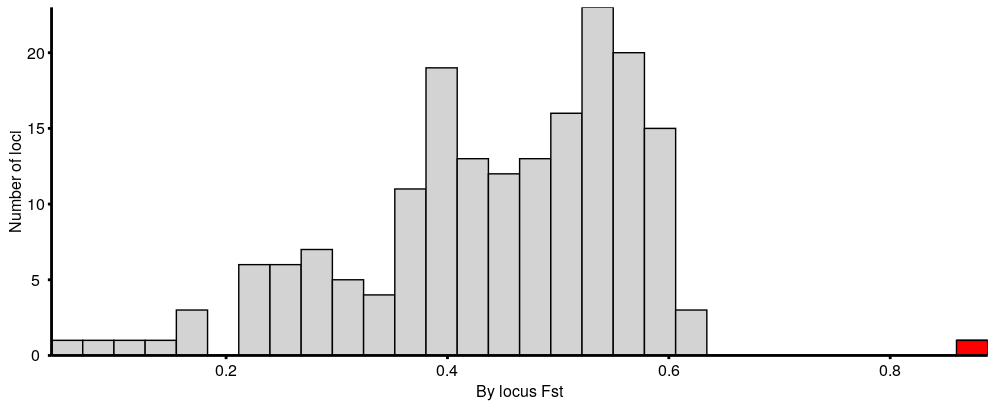

**Fig. S10:** Distribution of the 181 monolocus global *F_ST_* among the three populations of *Venturia inaequalis* defined by their host populations (*Malus domestica*, *M. sieversii* and hybrids). The red bar represents the global *F_ST_* estimated at the *V_081690_319* locus, which is the outlier in Fig. S9.

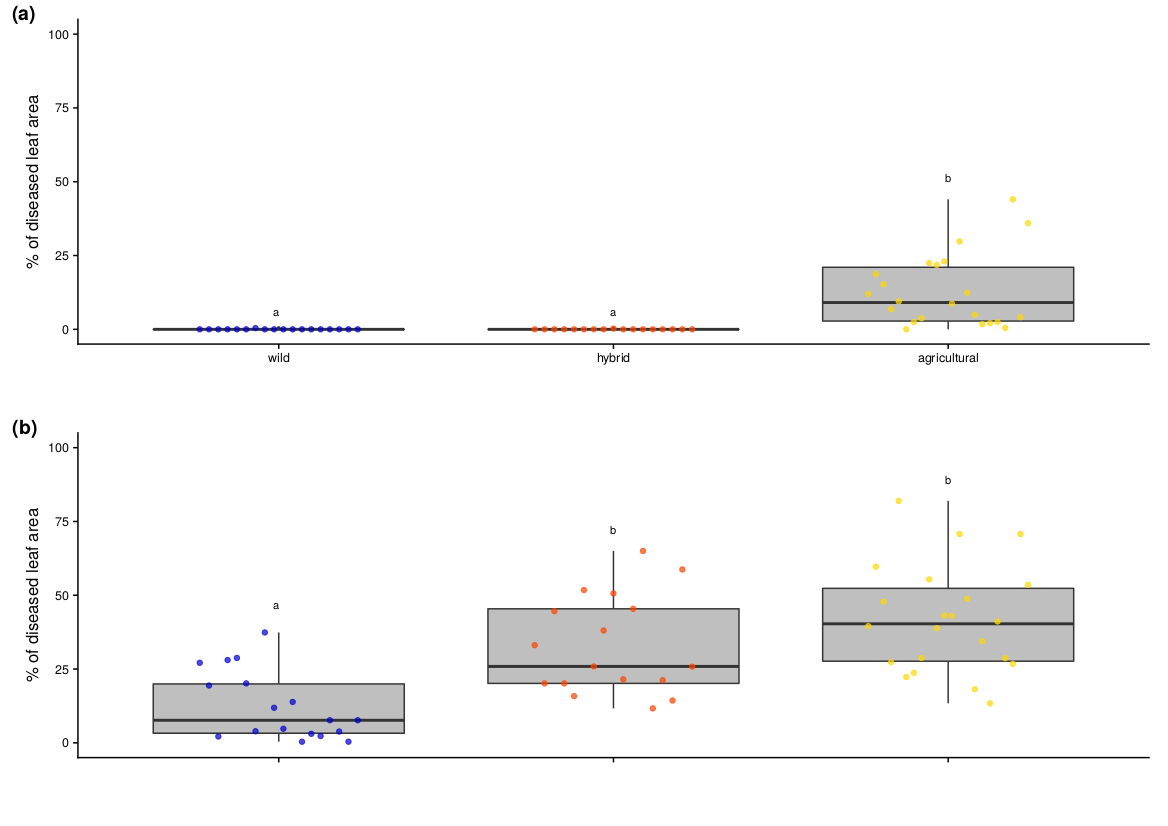

**Figure S11:** Results of pathogenicity experiments in controlled conditions with artificial inoculation of the fungus *Venturia inaequalis* (57 fungal strains; mean of three replicates per strain, one genotype per apple species) on apple tree leaves (*Malus sieversii* and *M. domestica*). This figure is the same as Figure 6, but with ancestry determined with the same threshold as for apple trees (wild type-type: *P_agr_*<0.2; hybrids: 0.2$\leq$*P_agr_*$\leq$0.8; agricultural: *P_agr_*>0.8). Boxplots of the percentage of scabbed leaf area for wild-type, agricultural-type and hybrid *V. inaequalis* strains, on A) *M. domestica* at 21 days post inoculation (dpi) and B) *M. sieversii* at 19 dpi. The box represents the lower and the upper quartiles. The thick horizontal line represents the median. The whiskers represent the largest and lowest observed values that fall within the distance of 1.5 times the interquartile range. The points represent the mean values of the percentage of diseased leaf area for each strain across the three replicates. Different letters indicate significant differences between populations (*P*<0.05; Wilcoxon’s rank sum tests).

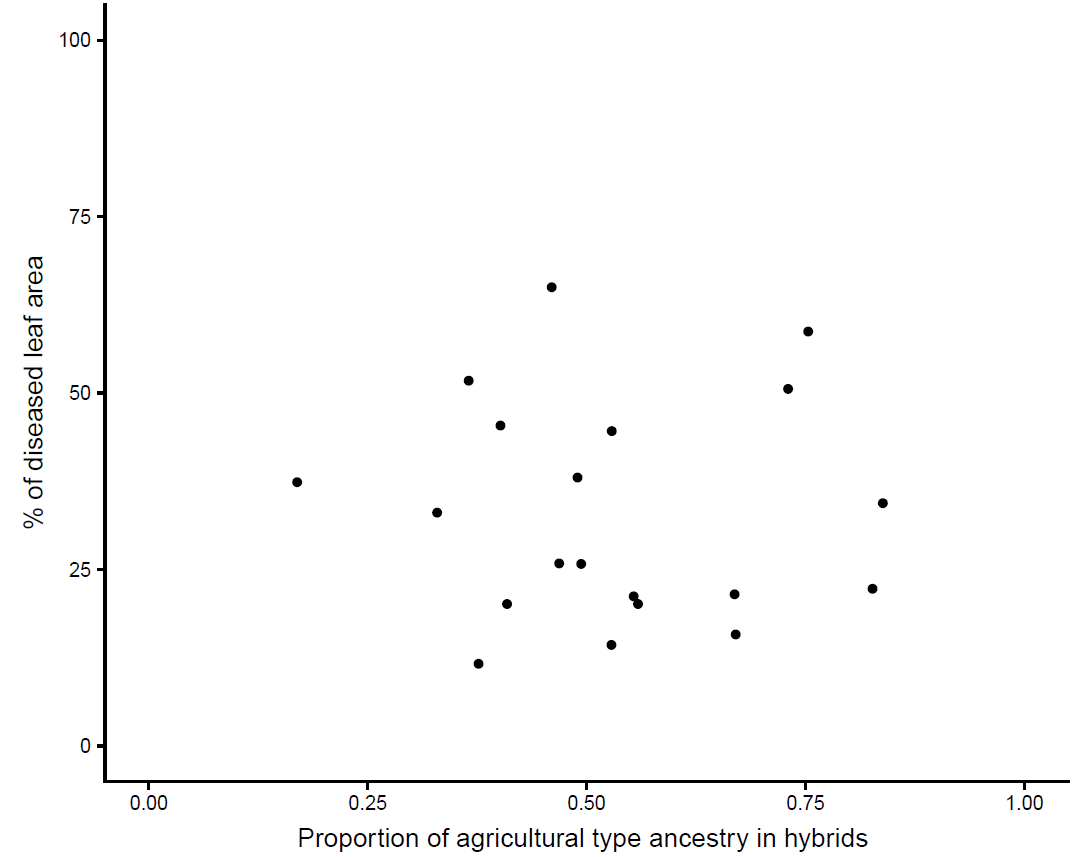

**Figure S12:** Percentage of scabbed *Malus sieversii* leaf area at 19 dpi plotted against the proportion of agricultural type ancestry in the *Venturia inaequalis* hybrid fungal strains.

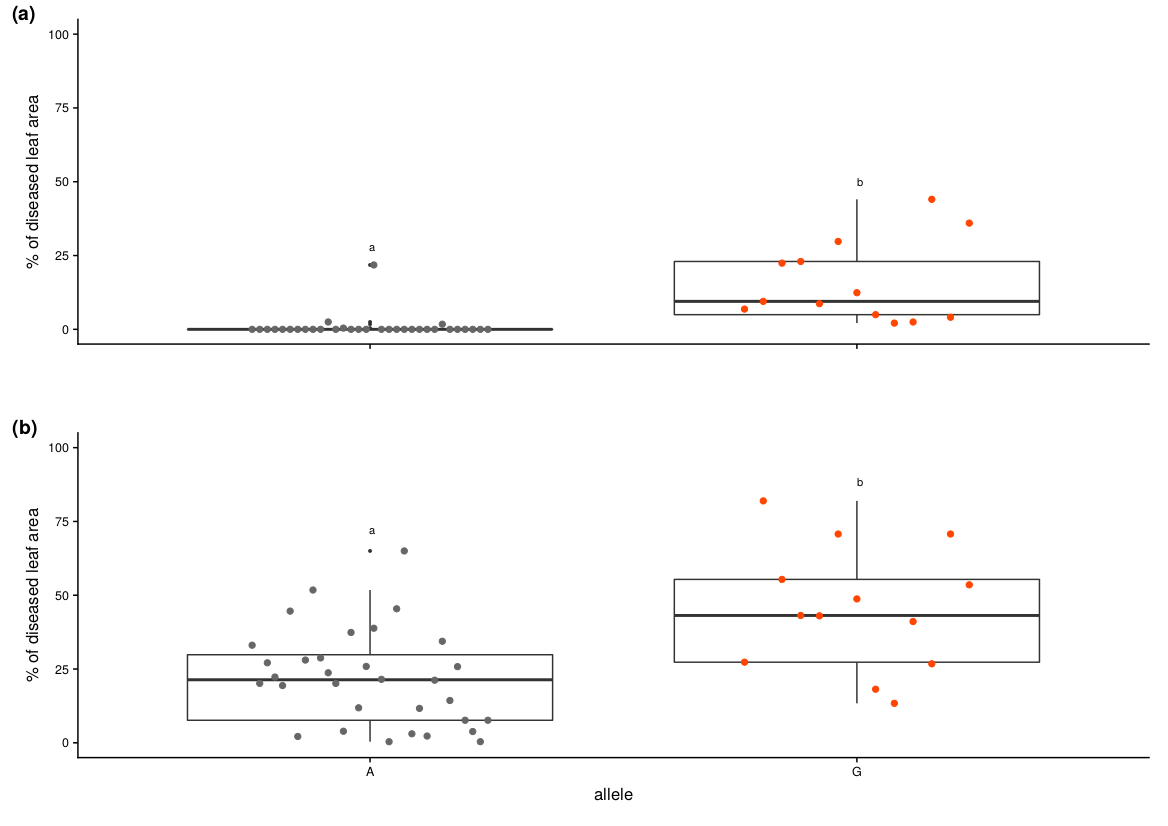

**Figure S13:** Results of pathogenicity experiments in controlled conditions with artificial inoculation of the fungus *Venturia inaequalis* (47 fungal strains; mean of three replicates per strain, one genotype per apple species) on apple tree leaves of *M. domestica* at 21 days post inoculation (dpi) (A) or *Malus sieversii* at 19 dpi (B). Boxplots of the percentage of scabbed leaf area for *V. inaequalis* strains carrying the A allele (TAC codon, corresponding to a tyrosine) and the G allele (TGA codon, corresponding to a codon stop) at the SNP *V_081690_319*. The box represents the lower and the upper quartiles. The thick horizontal line represents the median. The whiskers represent the largest and lowest observed values that fall within the distance of 1.5 times the interquartile range. The points represent the mean values of the percentage of diseased leaf area for each strain across the three replicates. Different letters indicate significant differences between populations (*P*<0.05; Wilcoxon’s rank sum tests).

**Table S1:** Description of *Venturia inaequalis* strains and *Malus spp.* trees sampled in 2012 in the Kazakh Tian Shan Mountains. Codes of the sampling locations: kot at Koturbulac; tag and tar at Talghar; lac, gec and esi at Esik; tuk and tum at Turgen.

P_agr_ : proportion of ancestry in the agricultural-type population of *V. inaequalis*

P_dom_ : proportion of ancestry in the *M. domestica* genepool

- Strain or tree not kept in the analysis because of missing data

* Strain used in pathological tests in controlled conditions

| Sampling site | Site description | Longitude | Latitude | Sample | Tree | Tree species | Number of microsatellites | P_dom_ | Strain | Fungal type | Number of SNP | P_agr_ |
| --- | --- | --- | --- | --- | --- | --- | --- | --- | --- | --- | --- | --- |
| esi_f | forest | 77.5253 | 43.31279 | fruit | 1 | M. sieversii | 26 | 0 | 12KZ307 | wild | 180 | 0 |
| esi_f | forest | 77.5253 | 43.31279 | fruit | 1 | M. sieversii | 26 | 0 | 12KZ328 | wild | 179 | 0.0284558 |
| esi_f | forest | 77.52547 | 43.31293 | leaf | 2 | M. sieversii | 25 | 0.0585092 | 12KZ308 | wild | 180 | 0.0559894 |
| esi_f | forest | 77.52547 | 43.31293 | leaf | 2 | M. sieversii | 25 | 0.0585092 | 12KZ329 | wild | 175 | 0.0420424 |
| esi_f | forest | 77.52668 | 43.31322 | leaf | 3 | M. sieversii | 25 | 0.0318404 | 12KZ309 | wild | 178 | 0.036804 |
| esi_f | forest | 77.52668 | 43.31322 | leaf | 3 | M. sieversii | 25 | 0.0318404 | 12KZ330 | agricultural | 175 | 0.9038027 |
| esi_f | forest | 77.52682 | 43.31319 | leaf | 4 | M. sieversii | 26 | 0.0788401 | 12KZ310 | wild | 171 | 0.0303849 |
| esi_f | forest | 77.52682 | 43.31319 | fruit | 4 | M. sieversii | 26 | 0.0788401 | 12KZ331 | wild | 178 | 0.0243452 |
| esi_f | forest | 77.52707 | 43.31326 | leaf | 5 | M. sieversii | 27 | 0 | 12KZ311 | agricultural | 178 | 0.9442636 |
| esi_f | forest | 77.52707 | 43.31326 | leaf | 5 | M. sieversii | 27 | 0 | 12KZ332 | wild | 175 | 0.024432 |
| esi_f | forest | 77.52715 | 43.31334 | leaf | 6 | M. sieversii | 20 | 0 | 12KZ333 | hybrid | 178 | 0.1643838 |
| esi_f | forest | 77.52722 | 43.31355 | leaf | 7 | M. sieversii | 25 | 0 | 12KZ313 | agricultural | 175 | 0.9354313 |
| esi_f | forest | 77.52721 | 43.31358 | leaf | 8 | M. sieversii | 26 | 0 | 12KZ314* | wild | 170 | 0 |
| esi_f | forest | 77.52721 | 43.31358 | leaf | 8 | M. sieversii | 26 | 0 | 12KZ335 | agricultural | 177 | 0.9613074 |
| esi_f | forest | 77.52763 | 43.3136 | leaf | 9 | M. sieversii | 25 | 0.1344876 | 12KZ315 | agricultural | 177 | 1 |
| esi_f | forest | 77.52769 | 43.31353 | leaf | 10 | M. sieversii | 27 | 0 | - | - |  | - |
| esi_f | forest | 77.52768 | 43.31351 | leaf | 11 | M. sieversii | 26 | 0 | 12KZ317 | agricultural | 176 | 0.9405866 |
| esi_f | forest | 77.52769 | 43.31345 | leaf | 12 | M. sieversii | 26 | 0.1858637 | 12KZ318 | agricultural | 176 | 0.9156555 |
| esi_f | forest | 77.52769 | 43.31345 | leaf | 12 | M. sieversii | 26 | 0.1858637 | 12KZ337 | agricultural | 175 | 0.9569234 |
| esi_f | forest | 77.52768 | 43.31345 | leaf | 13 | hybrid | 27 | 0.3779515 | 12KZ319 | agricultural | 161 | 0.9731225 |
| esi_f | forest | 77.52768 | 43.31345 | leaf | 13 | hybrid | 27 | 0.3779515 | 12KZ338* | agricultural | 177 | 0.9799088 |
| esi_f | forest | 77.52787 | 43.31341 | leaf | 14 | M. sieversii | 26 | 0 | 12KZ320 | agricultural | 178 | 0.9765746 |
| esi_f | forest | 77.52818 | 43.31348 | leaf | 15 | hybrid | 27 | 0.2734138 | 12KZ321* | agricultural | 176 | 0.9820073 |
| esi_f | forest | 77.52828 | 43.31367 | leaf | 16 | hybrid | 26 | 0.4176344 | 12KZ322 | agricultural | 175 | 0.9816884 |
| esi_f | forest | 77.52828 | 43.31367 | leaf | 16 | hybrid | 26 | 0.4176344 | 12KZ340 | agricultural | 178 | 0.9618765 |
| esi_f | forest | 77.52833 | 43.31368 | leaf | 17 | M. sieversii | 26 | 0 | - | - |  | - |
| esi_f | forest | 77.52841 | 43.31382 | leaf | 18 | hybrid | 27 | 0.4731279 | - | - |  | - |
| esi_f | forest | 77.52807 | 43.31364 | leaf | 19 | M. sieversii | 23 | 0.1129347 | 12KZ325* | agricultural | 177 | 0.9733975 |
| esi_f | forest | 77.52807 | 43.31364 | leaf | 19 | M. sieversii | 23 | 0.1129347 | 12KZ341 | agricultural | 177 | 1 |
| esi_f | forest | 77.52789 | 43.31373 | leaf | 20 | M. sieversii | 22 | 0.1362781 | 12KZ326 | agricultural | 179 | 0.9700057 |
| esi_f | forest | 77.52789 | 43.31373 | leaf | 20 | M. sieversii | 22 | 0.1362781 | 12KZ342 | agricultural | 178 | 0.9512272 |
| esi_f | forest | 77.52769 | 43.3137 | leaf | 21 | hybrid | 24 | 0.6526572 | 12KZ327 | agricultural | 178 | 0.9906453 |
| esi_f | forest | 77.52769 | 43.3137 | leaf | 21 | hybrid | 24 | 0.6526572 | 12KZ343* | agricultural | 178 | 0.9832528 |
| esi_f | forest | 77.52785 | 43.31353 | leaf | 22 | hybrid | 26 | 0.5810825 | - | - |  | - |
| esi_o | orchard | 77.52276 | 43.31351 | leaf | 1 | M. domestica | 27 | 0.9409015 | 12KZ081 | agricultural | 176 | 0.9559649 |
| esi_o | orchard | 77.52528 | 43.31344 | leaf | 2 | M. domestica | 27 | 1 | 12KZ082 | agricultural | 177 | 0.9721234 |
| esi_o | orchard | 77.52522 | 43.31313 | leaf | 3 | M. domestica | 26 | 1 | 12KZ092 | agricultural | 178 | 0.978869 |
| esi_o | orchard | 77.52558 | 43.31316 | leaf | 4 | M. domestica | 27 | 0.968369 | 12KZ084 | agricultural | 177 | 0.9475396 |
| esi_o | orchard | 77.52558 | 43.31316 | leaf | 4 | M. domestica | 27 | 0.968369 | 12KZ127 | agricultural | 175 | 0.9466059 |
| esi_o | orchard | 77.52536 | 43.31341 | leaf | 5 | M. domestica | 28 | 1 | 12KZ085 | agricultural | 176 | 0.9544723 |
| esi_o | orchard | 77.52536 | 43.31341 | leaf | 5 | M. domestica | 28 | 1 | 12KZ114 | agricultural | 168 | 0.904004 |
| esi_o | orchard | 77.52522 | 43.31339 | leaf | 6 | M. domestica | 27 | 1 | 12KZ086 | agricultural | 179 | 0.9004971 |
| esi_o | orchard | 77.52557 | 43.3135 | leaf | 7 | M. domestica | 28 | 1 | 12KZ087 | agricultural | 177 | 0.9665491 |
| esi_o | orchard | 77.52602 | 43.31352 | leaf | 8 | M. domestica | 24 | 0.815636 | 12KZ088 | agricultural | 178 | 0.9822238 |
| esi_o | orchard | 77.52687 | 43.31344 | leaf | 9 | M. domestica | 24 | 0.8944586 | 12KZ089 | agricultural | 178 | 0.9757034 |
| esi_o | orchard | 77.52687 | 43.31344 | leaf | 9 | M. domestica | 24 | 0.8944586 | 12KZ116 | agricultural | 176 | 0.9749681 |
| esi_o | orchard | 77.52528 | 43.31297 | leaf | 10 | M. domestica | 27 | 1 | 12KZ090* | agricultural | 180 | 0.9711211 |
| esi_o | orchard | 77.52625 | 43.31351 | leaf | 11 | M. domestica | 25 | 1 | 12KZ117 | agricultural | 175 | 0.9523531 |
| esi_o | orchard | 77.52663 | 43.31345 | leaf | 12 | M. domestica | 24 | 1 | 12KZ094* | agricultural | 177 | 1 |
| esi_o | orchard | 77.52639 | 43.31354 | leaf | 13 | M. domestica | 28 | 0.9089985 | 12KZ095 | agricultural | 171 | 0.9383283 |
| esi_o | orchard | 77.52682 | 43.31345 | leaf | 14 | M. domestica | 26 | 0.8894488 | - | - |  | - |
| esi_o | orchard | 77.52671 | 43.3133 | leaf | 15 | M. domestica | 28 | 0.8354614 | 12KZ097 | agricultural | 176 | 0.9729808 |
| esi_o | orchard | 77.52663 | 43.3133 | leaf | 16 | M. domestica | 28 | 0.8133564 | - | - |  | - |
| esi_o | orchard | 77.52654 | 43.31327 | leaf | 17 | M. domestica | 27 | 0.8899718 | 12KZ099 | agricultural | 175 | 0.9572485 |
| esi_o | orchard | 77.52654 | 43.31327 | leaf | 17 | M. domestica | 27 | 0.8899718 | 12KZ119 | agricultural | 178 | 0.9820246 |
| esi_o | orchard | 77.52618 | 43.31323 | leaf | 18 | M. domestica | 28 | 0.8354614 | 12KZ100 | agricultural | 179 | 0.9604833 |
| esi_o | orchard | 77.52595 | 43.31348 | leaf | 19 | M. domestica | 27 | 0.9196755 | 12KZ121* | agricultural | 176 | 0.9913167 |
| esi_o | orchard | 77.5264 | 43.31324 | leaf | 20 | M. domestica | 27 | 0.8261757 | 12KZ120 | agricultural | 176 | 0.9533324 |
| esi_o | orchard | 77.52541 | 43.31315 | leaf | 21 | M. domestica | 27 | 0.9714831 | - | - |  | - |
| esi_o | orchard | 77.52595 | 43.31322 | leaf | 22 | M. domestica | 26 | 0.8878602 | 12KZ123 | agricultural | 179 | 0.9860752 |
| esi_o | orchard | 77.5258 | 43.31316 | leaf | 23 | M. domestica | 27 | 0.9634318 | 12KZ105 | agricultural | 178 | 0.9720818 |
| esi_o | orchard | 77.5258 | 43.31316 | leaf | 23 | M. domestica | 27 | 0.9634318 | 12KZ124 | agricultural | 164 | 0.9463758 |
| esi_o | orchard | 77.52608 | 43.31317 | leaf | 24 | M. domestica | 28 | 0.8354614 | 12KZ106 | agricultural | 178 | 0.9725201 |
| esi_o | orchard | 77.52608 | 43.31317 | leaf | 24 | M. domestica | 28 | 0.8354614 | 12KZ125* | agricultural | 172 | 1 |
| esi_o | orchard | 77.5265 | 43.31355 | leaf | 25 | M. domestica | 25 | 0.972823 | - | - |  | - |
| esi_o | orchard | 77.52695 | 43.31322 | leaf | 26 | hybrid | 26 | 0.5550558 | 12KZ108 | agricultural | 175 | 0.9667686 |
| esi_o | orchard | 77.52553 | 43.31343 | leaf | 27 | M. domestica | 28 | 0.9728445 | 12KZ109 | agricultural | 174 | 0.9869203 |
| esi_o | orchard | 77.52687 | 43.31339 | leaf | 28 | M. domestica | 27 | 1 | - | - |  | - |
| esi_o | orchard | 77.52699 | 43.31344 | leaf | 29 | M. domestica | 24 | 0.889702 | 12KZ111 | agricultural | 175 | 0.9461959 |
| esi_o | orchard | 77.52698 | 43.31329 | leaf | 30 | hybrid | 26 | 0.7916631 | - | - |  | - |
| gec | forest | 77.49844 | 43.27076 | leaf | 1 | M. sieversii | 28 | 0.0532503 | 12KZ181* | agricultural | 176 | 1 |
| gec | forest | 77.49831 | 43.27066 | leaf | 2 | M. sieversii | 27 | 0.0360634 | 12KZ182 | wild | 180 | 0.0243931 |
| gec | forest | 77.49816 | 43.27058 | leaf | 3 | M. sieversii | 28 | 0 | 12KZ183 | agricultural | 175 | 0.9254778 |
| gec | forest | 77.49747 | 43.27002 | leaf | 4 | M. sieversii | 28 | 0 | 12KZ184 | wild | 179 | 0 |
| gec | forest | 77.49705 | 43.2697 | leaf | 5 | M. sieversii | 28 | 0.0453792 | 12KZ185 | wild | 178 | 0.0232814 |
| gec | forest | 77.49708 | 43.26944 | leaf | 6 | M. sieversii | 28 | 0.066967 | 12KZ186 | agricultural | 178 | 0.949053 |
| gec | forest | 77.49856 | 43.27144 | leaf | 7 | M. sieversii | 25 | 0.0805205 | 12KZ187* | hybrid | 174 | 0.5284469 |
| gec | forest | 77.4986 | 43.27153 | leaf | 8 | M. sieversii | 25 | 0.0805205 | 12KZ188* | hybrid | 173 | 0.5587992 |
| gec | forest | 77.49866 | 43.27158 | leaf | 9 | M. sieversii | 26 | 0.0345664 | 12KZ189 | wild | 175 | 0.0412516 |
| gec | forest | 77.49857 | 43.27159 | leaf | 10 | M. sieversii | 25 | 0 | - | - |  | - |
| gec | forest | 77.49855 | 43.27158 | leaf | 11 | M. sieversii | 26 | 0.0749451 | 12KZ191* | hybrid | 175 | 0.7531344 |
| gec | forest | 77.49851 | 43.27157 | leaf | 12 | M. sieversii | 28 | 0.0657692 | 12KZ192 | wild | 176 | 0.0231789 |
| gec | forest | 77.49851 | 43.27157 | leaf | 12 | M. sieversii | 28 | 0.0657692 | 12KZ193 | agricultural | 176 | 1 |
| gec | forest | 77.4985 | 43.27156 | leaf | 13 | M. sieversii | 27 | 0.0713742 | 12KZ194* | hybrid | 175 | 0.4938612 |
| gec | forest | 77.4985 | 43.27156 | leaf | 13 | M. sieversii | 27 | 0.0713742 | 12KZ195* | hybrid | 179 | 0.6703498 |
| gec | forest | 77.49849 | 43.27153 | leaf | 14 | M. sieversii | 27 | 0.0833405 | - | - | - | - |
| gec | forest | 77.49849 | 43.27153 | leaf | 14 | M. sieversii | 27 | 0.0833405 | 12KZ197 | hybrid | 179 | 0.2105954 |
| gec | forest | 77.49845 | 43.27161 | leaf | 15 | M. sieversii | 28 | 0.0484277 | 12KZ198 | agricultural | 153 | 0.9854747 |
| gec | forest | 77.49845 | 43.27161 | leaf | 15 | M. sieversii | 28 | 0.0484277 | 12KZ199 | agricultural | 161 | 0.9282926 |
| gec | forest | 77.49834 | 43.27164 | leaf | 16 | M. sieversii | 27 | 0.0404232 | - | - | - | - |
| gec | forest | 77.49831 | 43.27164 | leaf | 17 | M. sieversii | 27 | 0 | 12KZ165* | hybrid | 178 | 0.8267053 |
| gec | forest | 77.49831 | 43.27164 | leaf | 17 | M. sieversii | 27 | 0 | 12KZ166 | agricultural | 174 | 0.9722035 |
| gec | forest | 77.49936 | 43.27172 | leaf | 18 | M. sieversii | 27 | 0.0523775 | 12KZ167* | hybrid | 178 | 0.5538649 |
| gec | forest | 77.49931 | 43.2718 | leaf | 19 | M. sieversii | 28 | 0.0361204 | - | - |  | - |
| gec | forest | 77.49856 | 43.27188 | leaf | 20 | M. sieversii | 27 | 0.1288788 | - | - |  | - |
| gec | forest | 77.49863 | 43.27195 | leaf | 21 | M. sieversii | 22 | 0.039094 | 12KZ170* | wild | 176 | 0 |
| gec | forest | 77.49866 | 43.27188 | leaf | 22 | M. sieversii | 20 | 0.1416511 | 12KZ171 | wild | 177 | 0.0251143 |
| gec | forest | 77.49869 | 43.27184 | leaf | 23 | M. sieversii | 22 | 0.128774 | 12KZ172 | wild | 176 | 0.0517918 |
| gec | forest | 77.49878 | 43.2719 | leaf | 24 | M. sieversii | 18 | 0 | 12KZ173* | hybrid | 178 | 0.5288262 |
| gec | forest | 77.49879 | 43.27188 | leaf | 25 | M. sieversii | 19 | 0 | - | - |  | - |
| gec | forest | 77.49883 | 43.27189 | leaf | 26 | M. sieversii | 21 | 0.0388755 | - | - |  | - |
| gec | forest | 77.49888 | 43.2719 | leaf | 27 | M. sieversii | 23 | 0.0362793 | 12KZ175 | wild | 179 | 0.0212231 |
| gec | forest | 77.49889 | 43.27188 | leaf | 28 | M. sieversii | 25 | 0.0362228 | - | - |  | - |
| gec | forest | 77.49887 | 43.2718 | leaf | 29 | M. sieversii | 26 | 0.0299394 | 12KZ177* | hybrid | 180 | 0.8384605 |
| gec | forest | 77.49866 | 43.27181 | leaf | 30 | M. sieversii | 28 | 0.1402292 | - | - |  | - |
| gec | forest | 77.49832 | 43.27184 | leaf | 31 | M. sieversii | 23 | 0.1089276 | 12KZ179 | wild | 179 | 0.0137093 |
| kot | forest | 77.06929 | 43.25708 | leaf | 1 | M. sieversii | 26 | 0.0380377 | 12KZ016 | wild | 181 | 0.014037 |
| kot | forest | 77.06929 | 43.25708 | leaf | 1 | M. sieversii | 26 | 0.0380377 | 12KZ038 | wild | 180 | 0.0297141 |
| kot | forest | 77.06935 | 43.25711 | leaf | 2 | M. sieversii | 27 | 0 | 12KZ011 | wild | 180 | 0.0236077 |
| kot | forest | 77.06935 | 43.25711 | leaf | 2 | M. sieversii | 27 | 0 | 12KZ028 | wild | 179 | 0.0208921 |
| kot | forest | 77.0694 | 43.25711 | leaf | 3 | M. sieversii | 28 | 0.0346958 | 12KZ006 | wild | 180 | 0.0350002 |
| kot | forest | 77.0694 | 43.25711 | leaf | 3 | M. sieversii | 28 | 0.0346958 | 12KZ023 | wild | 178 | 0.0269538 |
| kot | forest | 77.06946 | 43.25706 | leaf | 4 | M. sieversii | 28 | 0.0288201 | 12KZ007 | wild | 180 | 0.0248488 |
| kot | forest | 77.06915 | 43.25714 | leaf | 5 | hybrid | 28 | 0.7187674 | 12KZ019 | agricultural | 178 | 0.9382779 |
| kot | forest | 77.06915 | 43.25714 | leaf | 5 | hybrid | 28 | 0.7187674 | 12KZ034 | agricultural | 165 | 0.9763383 |
| kot | forest | 77.06924 | 43.2572 | leaf | 6 | M. sieversii | 28 | 0.1241035 | 12KZ001 | wild | 177 | 0 |
| kot | forest | 77.06924 | 43.25727 | leaf | 7 | M. sieversii | 26 | 0 | 12KZ008* | agricultural | 177 | 0.9789762 |
| kot | forest | 77.06679 | 43.25947 | leaf | 8 | - | - | - | 12KZ022* | agricultural | 178 | 0.9781252 |
| kot | forest | 77.06679 | 43.25947 | leaf | 8 | - | - | - | 12KZ029* | hybrid | 180 | 0.7302151 |
| kot | forest | 77.06767 | 43.25971 | leaf | 9 | hybrid | 26 | 0.3486668 | 12KZ014 | agricultural | 179 | 0.9277547 |
| kot | forest | 77.06767 | 43.25971 | leaf | 9 | hybrid | 26 | 0.3486668 | 12KZ032 | agricultural | 178 | 1 |
| kot | forest | 77.06761 | 43.25983 | leaf | 10 | M. sieversii | 28 | 0.0436823 | 12KZ003 | wild | 175 | 0.0203183 |
| kot | forest | 77.06761 | 43.25983 | leaf | 10 | M. sieversii | 28 | 0.0436823 | 12KZ020 | wild | 181 | 0.040354 |
| kot | forest | 77.06761 | 43.25983 | leaf | 10 | M. sieversii | 28 | 0.0436823 | 12KZ042 | wild | 179 | 0.0334561 |
| kot | forest | 77.06755 | 43.2598 | leaf | 11 | M. sieversii | 25 | 0 | - | - |  | - |
| kot | forest | 77.06775 | 43.26015 | leaf | 12 | M. sieversii | 25 | 0.0861022 | 12KZ009 | agricultural | 178 | 0.9466606 |
| kot | forest | 77.06775 | 43.26015 | leaf | 12 | M. sieversii | 25 | 0.0861022 | 12KZ026* | hybrid | 178 | 0.3293489 |
| kot | forest | 77.06775 | 43.26015 | leaf | 12 | M. sieversii | 25 | 0.0861022 | 12KZ035 | agricultural | 177 | 0.9872486 |
| kot | forest | 77.06732 | 43.26026 | leaf | 13 | M. sieversii | 22 | 0.1923703 | 12KZ012* | agricultural | 179 | 0.9710179 |
| kot | forest | 77.06732 | 43.26026 | leaf | 13 | M. sieversii | 22 | 0.1923703 | 12KZ030* | agricultural | 175 | 0.9844703 |
| kot | forest | 77.06732 | 43.26026 | leaf | 13 | M. sieversii | 22 | 0.1923703 | 12KZ044 | agricultural | 179 | 0.9764536 |
| kot | forest | 77.06729 | 43.26034 | leaf | 14 | M. sieversii | 27 | 0.0825608 | 12KZ017 | wild | 177 | 0.0483149 |
| kot | forest | 77.0668 | 43.26058 | leaf | 15 | hybrid | 20 | 0.2203774 | 12KZ033* | hybrid | 179 | 0.6691175 |
| kot | forest | 77.0668 | 43.26058 | leaf | 15 | hybrid | 20 | 0.2203774 | 12KZ037 | agricultural | 177 | 0.940881 |
| kot | forest | 77.06652 | 43.26067 | leaf | 16 | M. sieversii | 26 | 0 | 12KZ010 | wild | 178 | 0.0172472 |
| kot | forest | 77.06579 | 43.26113 | leaf | 17 | hybrid | 26 | 0.5550558 | - | - | - | - |
| kot | forest | 77.06579 | 43.26113 | leaf | 17 | hybrid | 26 | 0.5550558 | 12KZ036* | agricultural | 178 | 1 |
| lac | forest | 77.48412 | 43.25617 | leaf | 1 | M. sieversii | 25 | 0 | 12KZ130* | wild | 180 | 0.0126477 |
| lac | forest | 77.48401 | 43.25615 | leaf | 2 | M. sieversii | 28 | 0 | 12KZ131 | wild | 178 | 0.0346555 |
| lac | forest | 77.48401 | 43.25615 | leaf | 2 | M. sieversii | 28 | 0 | 12KZ132 | wild | 169 | 0.0166933 |
| lac | forest | 77.48396 | 43.25618 | leaf | 3 | M. sieversii | 28 | 0.0358908 | 12KZ134 | wild | 180 | 0.0345627 |
| lac | forest | 77.484 | 43.25647 | leaf | 4 | M. sieversii | 27 | 0 | 12KZ135 | wild | 177 | 0.0262321 |
| lac | forest | 77.484 | 43.25647 | leaf | 4 | M. sieversii | 27 | 0 | 12KZ136 | wild | 168 | 0.0273367 |
| lac | forest | 77.4844 | 43.257 | leaf | 5 | M. sieversii | 25 | 0.0362969 | 12KZ137 | wild | 179 | 0.0387558 |
| lac | forest | 77.4844 | 43.257 | leaf | 5 | M. sieversii | 25 | 0.0362969 | 12KZ162 | wild | 180 | 0.029449 |
| lac | forest | 77.48494 | 43.25685 | leaf | 6 | M. sieversii | 28 | 0.0456372 | 12KZ139 | wild | 177 | 0.0354743 |
| lac | forest | 77.48494 | 43.25685 | leaf | 6 | M. sieversii | 28 | 0.0456372 | 12KZ140 | wild | 177 | 0.0302848 |
| lac | forest | 77.48494 | 43.25685 | leaf | 6 | M. sieversii | 28 | 0.0456372 | 12KZ163 | agricultural | 178 | 0.9508097 |
| lac | forest | 77.48494 | 43.25685 | leaf | 6 | M. sieversii | 28 | 0.0456372 | 12KZ164 | wild | 176 | 0.0127806 |
| lac | forest | 77.4849 | 43.25674 | leaf | 7 | M. sieversii | 27 | 0.0424196 | 12KZ141* | wild | 179 | 0 |
| lac | forest | 77.4849 | 43.25674 | leaf | 7 | M. sieversii | 27 | 0.0424196 | 12KZ142 | wild | 175 | 0 |
| lac | forest | 77.48514 | 43.25673 | leaf | 8 | M. sieversii | 28 | 0 | 12KZ144 | wild | 181 | 0.045821 |
| lac | forest | 77.48514 | 43.25673 | leaf | 8 | M. sieversii | 28 | 0 | 12KZ146* | hybrid | 154 | 0.3766908 |
| lac | forest | 77.48535 | 43.25674 | leaf | 9 | M. domestica | 28 | 1 | 12KZ147 | agricultural | 176 | 0.9198747 |
| lac | forest | 77.48535 | 43.25674 | leaf | 9 | M. domestica | 28 | 1 | 12KZ148 | agricultural | 178 | 0.9370156 |
| lac | forest | 77.48535 | 43.25674 | leaf | 9 | M. domestica | 28 | 1 | 12KZ149 | agricultural | 178 | 0.9519796 |
| lac | forest | 77.4854 | 43.25681 | leaf | 10 | M. sieversii | 26 | 0.1651177 | 12KZ150 | agricultural | 171 | 0.9574846 |
| lac | forest | 77.4854 | 43.25681 | leaf | 10 | M. sieversii | 26 | 0.1651177 | 12KZ151 | agricultural | 172 | 0.9473876 |
| lac | forest | 77.48553 | 43.25681 | leaf | 11 | hybrid | 26 | 0.3235387 | - | - | - | - |
| lac | forest | 77.48553 | 43.25681 | leaf | 11 | hybrid | 26 | 0.3235387 | 12KZ152 | agricultural | 179 | 0.9618752 |
| lac | forest | 77.48563 | 43.25699 | leaf | 12 | M. sieversii | 26 | 0.0672049 | 12KZ155 | wild | 180 | 0.025655 |
| lac | forest | 77.48563 | 43.25699 | leaf | 12 | M. sieversii | 26 | 0.0672049 | 12KZ161 | wild | 174 | 0.0265095 |
| lac | forest | 77.48489 | 43.2565 | leaf | 13 | M. sieversii | 28 | 0.0873071 | 12KZ156* | wild | 178 | 0.0091008 |
| lac | forest | 77.48489 | 43.2565 | leaf | 13 | M. sieversii | 28 | 0.0873071 | 12KZ157 | wild | 178 | 0.0192818 |
| lac | forest | 77.48489 | 43.2565 | leaf | 13 | M. sieversii | 28 | 0.0873071 | 12KZ160 | wild | 180 | 0 |
| lac | forest | 77.4815 | 43.25605 | leaf | 14 | hybrid | 27 | 0.5166333 | 12KZ158 | agricultural | 177 | 0.9705186 |
| lac | forest | 77.4815 | 43.25605 | leaf | 14 | hybrid | 27 | 0.5166333 | 12KZ159 | agricultural | 177 | 0.960681 |
| lac | forest | 77.48171 | 43.25599 | leaf | 15 | M. sieversii | 27 | 0.1708755 | - | - |  | - |
| lac | forest | 77.48171 | 43.25602 | leaf | 16 | hybrid | 28 | 0.5279454 | - | - |  | - |
| tag | forest | 77.21798 | 43.23539 | fruit | 1 | M. sieversii | 26 | 0.0583075 | 12KZ202* | wild | 179 | 0 |
| tag | forest | 77.21786 | 43.23565 | fruit | 2 | M. sieversii | 27 | 0.1009104 | 12KZ203* | wild | 178 | 0 |
| tag | forest | 77.21786 | 43.23565 | leaf | 2 | M. sieversii | 27 | 0.1009104 | 12KZ204 | agricultural | 177 | 0.9893188 |
| tag | forest | 77.21786 | 43.23535 | fruit | 3 | M. sieversii | 27 | 0.0614618 | 12KZ205 | wild | 169 | 0.0611573 |
| tag | forest | 77.21773 | 43.23619 | fruit | 4 | M. sieversii | 26 | 0.1458155 | 12KZ206 | agricultural | 171 | 1 |
| tag | forest | 77.21773 | 43.23619 | leaf | 4 | M. sieversii | 26 | 0.1458155 | 12KZ207* | agricultural | 178 | 0.9640103 |
| tag | forest | 77.21796 | 43.23541 | leaf | 5 | M. sieversii | 26 | 0.1458155 | 12KZ208 | agricultural | 176 | 0.9814613 |
| tag | forest | 77.21799 | 43.23569 | leaf | 6 | M. sieversii | 28 | 0.1322566 | - | - | - | - |
| tag | forest | 77.21802 | 43.23526 | leaf | 7 | M. sieversii | 28 | 0.0626317 | - | - |  | - |
| tag | forest | 77.21821 | 43.23569 | leaf | 8 | M. sieversii | 26 | 0.0704455 | 12KZ212* | wild | 180 | 0.0181697 |
| tag | forest | 77.21819 | 43.23564 | leaf | 9 | M. sieversii | 26 | 0 | 12KZ213 | wild | 177 | 0.0207722 |
| tag | forest | 77.21828 | 43.23531 | leaf | 10 | M. sieversii | 27 | 0 | 12KZ215 | wild | 166 | 0.0147369 |
| tag | forest | 77.21823 | 43.23563 | leaf | 11 | M. sieversii | 28 | 0 | 12KZ214* | hybrid | 176 | 0.3654056 |
| tag | forest | 77.21847 | 43.23557 | fruit | 12 | M. sieversii | 27 | 0 | - | - |  | - |
| tag | forest | 77.21839 | 43.23549 | leaf | 13 | M. sieversii | 28 | 0 | 12KZ217* | agricultural | 151 | 0.9887712 |
| tag | forest | 77.2185 | 43.23552 | leaf | 14 | M. sieversii | 28 | 0 | 12KZ219 | wild | 180 | 0.0264612 |
| tag | forest | 77.21844 | 43.23548 | leaf | 15 | M. sieversii | 27 | 0 | 12KZ218* | wild | 173 | 0 |
| tag | forest | 77.21791 | 43.23596 | fruit | 16 | M. sieversii | 27 | 0.0816537 | 12KZ220 | hybrid | 178 | 0.5178266 |
| tag | forest | 77.2178 | 43.23615 | leaf | 17 | M. sieversii | 27 | 0 | - | - | - | - |
| tag | forest | 77.2178 | 43.23615 | leaf | 18 | M. sieversii | 24 | 0 | 12KZ222 | wild | 164 | 0.0216918 |
| tag | forest | 77.21773 | 43.23624 | fruit | 19 | M. sieversii | 22 | 0.0496972 | 12KZ223 | wild | 180 | 0.0217218 |
| tag | forest | 77.21774 | 43.23622 | fruit | 20 | M. sieversii | 27 | 0 | - | - |  | - |
| tag | forest | 77.21825 | 43.23612 | fruit | 21 | M. sieversii | 28 | 0.1330408 | 12KZ225 | wild | 180 | 0.0131238 |
| tag | forest | 77.21825 | 43.23612 | leaf | 21 | M. sieversii | 28 | 0.1330408 | 12KZ226 | wild | 179 | 0 |
| tag | forest | 77.21828 | 43.23603 | leaf | 22 | M. sieversii | 27 | 0.0923417 | 12KZ227* | agricultural | 171 | 0.9780749 |
| tag | forest | 77.21845 | 43.23597 | fruit | 23 | hybrid | 27 | 0.3516571 | - | - | - | - |
| tag | forest | 77.21851 | 43.23602 | leaf | 24 | M. sieversii | 28 | 0.0346774 | 12KZ229 | hybrid | 176 | 0.366748 |
| tag | forest | 77.21858 | 43.23611 | leaf | 25 | hybrid | 23 | 0.2883515 | 12KZ230 | agricultural | 172 | 0.966946 |
| tag | forest | 77.21827 | 43.23675 | fruit | 26 | M. sieversii | 28 | 0.0451234 | 12KZ231* | agricultural | 177 | 0.9755135 |
| tag | forest | 77.21834 | 43.2364 | leaf | 27 | M. sieversii | 27 | 0.0872648 | 12KZ232* | wild | 180 | 0 |
| tag | forest | 77.21854 | 43.23629 | leaf | 28 | M. sieversii | 25 | 0.0452591 | 12KZ233 | agricultural | 170 | 0.988893 |
| tag | forest | 77.21877 | 43.23621 | fruit | 29 | M. sieversii | 28 | 0.0619082 | 12KZ235* | wild | 169 | 0.0131683 |
| tag | forest | 77.219 | 43.23629 | leaf | 30 | M. sieversii | 25 | 0.1049889 | 12KZ236* | wild | 179 | 0.0077767 |
| tag | forest | 77.21835 | 43.23671 | leaf | 31 | hybrid | 26 | 0.5211862 | - | - |  | - |
| tag | forest | 77.21814 | 43.23673 | leaf | 32 | hybrid | 25 | 0.5216352 | - | - |  | - |
| tag | forest | 77.2185 | 43.23642 | leaf | 33 | M. sieversii | 23 | 0 | - | - |  | - |
| tar | forest | 77.27885 | 43.23053 | fruit | 1 | hybrid | 25 | 0.304393 | 12KZ237* | agricultural | 177 | 0.9776292 |
| tar | forest | 77.27885 | 43.23053 | fruit | 1 | hybrid | 25 | 0.304393 | 12KZ354 | agricultural | 174 | 0.9765472 |
| tar | forest | 77.27921 | 43.23042 | fruit | 2 | M. sieversii | 25 | 0.1267311 | 12KZ303 | wild | 179 | 0.0406185 |
| tar | forest | 77.27917 | 43.23035 | fruit | 3 | hybrid | 22 | 0.3326421 | 12KZ242 | agricultural | 178 | 0.9525288 |
| tar | forest | 77.27906 | 43.2304 | fruit | 4 | M. sieversii | 27 | 0.1878046 | 12KZ244 | wild | 178 | 0.029532 |
| tar | forest | 77.27905 | 43.23034 | fruit | 5 | M. sieversii | 28 | 0 | 12KZ245 | wild | 179 | 0.0458792 |
| tar | forest | 77.27908 | 43.23014 | fruit | 6 | M. sieversii | 28 | 0.1012558 | - | - | - | - |
| tar | forest | 77.27913 | 43.23011 | fruit | 7 | M. sieversii | 26 | 0 | 12KZ247 | wild | 175 | 0.0346412 |
| tar | forest | 77.27946 | 43.23045 | leaf | 8 | hybrid | 27 | 0.2486991 | 12KZ304 | agricultural | 177 | 0.9791636 |
| tar | forest | 77.28102 | 43.22887 | fruit | 9 | hybrid | 22 | 0.7604932 | 12KZ249 | agricultural | 177 | 0.9633133 |
| tar | forest | 77.28113 | 43.22894 | fruit | 10 | M. sieversii | 25 | 0.0672512 | 12KZ250 | wild | 178 | 0.0135316 |
| tar | forest | 77.28122 | 43.22894 | fruit | 11 | M. sieversii | 28 | 0.0401198 | - | - | - | - |
| tar | forest | 77.2812 | 43.2289 | fruit | 12 | M. sieversii | 26 | 0 | 12KZ252 | hybrid | 177 | 0.493678 |
| tar | forest | 77.28129 | 43.22889 | leaf | 13 | M. sieversii | 28 | 0 | - | - | - | - |
| tar | forest | 77.28135 | 43.22886 | fruit | 14 | M. sieversii | 28 | 0.0305047 | 12KZ255 | wild | 180 | 0.0373872 |
| tar | forest | 77.28145 | 43.22888 | fruit | 15 | M. sieversii | 22 | 0 | 12KZ256* | hybrid | 164 | 0.1694653 |
| tar | forest | 77.28152 | 43.22877 | fruit | 16 | M. sieversii | 23 | 0.0769187 | - | - |  | - |
| tar | forest | 77.27956 | 43.22873 | fruit | 17 | M. sieversii | 27 | 0.0529684 | 12KZ258 | wild | 179 | 0.0699615 |
| tar | forest | 77.2816 | 43.22864 | fruit | 18 | M. sieversii | 23 | 0.1004873 | 12KZ259 | wild | 180 | 0.0714904 |
| tar | forest | 77.28163 | 43.22855 | fruit | 19 | M. sieversii | 24 | 0.0583591 | - | - | - | - |
| tar | forest | 77.28159 | 43.22855 | fruit | 20 | M. sieversii | 27 | 0.0463422 | - | - | - | - |
| tar | forest | 77.28172 | 43.22846 | fruit | 21 | M. sieversii | 23 | 0.046788 | 12KZ262 | wild | 171 | 0.0271832 |
| tar | forest | 77.28175 | 43.22847 | fruit | 22 | M. sieversii | 28 | 0 | 12KZ263 | hybrid | 174 | 0.100648 |
| tar | forest | 77.2804 | 43.22918 | fruit | 23 | M. sieversii | 26 | 0.050751 | 12KZ264 | wild | 179 | 0.0390532 |
| tar | forest | 77.28013 | 43.22933 | fruit | 24 | M. sieversii | 28 | 0 | 12KZ265 | wild | 178 | 0.0483447 |
| tar | forest | 77.28008 | 43.22932 | fruit | 25 | M. sieversii | 26 | 0 | 12KZ266* | wild | 179 | 0 |
| tar | forest | 77.27985 | 43.22938 | fruit | 26 | M. sieversii | 26 | 0.0514275 | 12KZ305 | wild | 174 | 0 |
| tar | forest | 77.27972 | 43.2295 | fruit | 27 | M. sieversii | 24 | 0.0823108 | 12KZ306* | hybrid | 177 | 0.4897067 |
| tar | forest | 77.27956 | 43.22982 | fruit | 28 | hybrid | 27 | 0.3238739 | 12KZ269* | agricultural | 157 | 0.9682965 |
| tar | forest | 77.27943 | 43.23021 | fruit | 29 | M. sieversii | 28 | 0.0576951 | 12KZ270 | agricultural | 174 | 0.9737557 |
| tar | forest | 77.27935 | 43.23038 | fruit | 30 | hybrid | 27 | 0.24589 | 12KZ271 | agricultural | 173 | 0.9736604 |
| tuk | forest | 77.67234 | 43.37067 | leaf | 1 | M. sieversii | 28 | 0 | 12KZ079 | wild | 180 | 0 |
| tuk | forest | 77.67196 | 43.37141 | leaf | 2 | M. sieversii | 28 | 0.041588 | 12KZ047 | wild | 164 | 0.0774264 |
| tuk | forest | 77.67207 | 43.37177 | leaf | 3 | M. sieversii | 28 | 0 | - | - |  | - |
| tuk | forest | 77.67184 | 43.37165 | leaf | 4 | M. sieversii | 22 | 0.1973146 | 12KZ048 | wild | 180 | 0.0152405 |
| tuk | forest | 77.67189 | 43.37156 | leaf | 5 | - | - | - | 12KZ049 | wild | 177 | 0.0235404 |
| tuk | forest | 77.67197 | 43.37166 | leaf | 6 | - | - | - | 12KZ050* | wild | 162 | 0.0126202 |
| tuk | forest | 77.67218 | 43.37159 | leaf | 7 | - | - | - | 12KZ051* | wild | 177 | 0 |
| tuk | forest | 77.67229 | 43.37111 | leaf | 8 | - | - | - | 12KZ052 | wild | 176 | 0.0325841 |
| tuk | forest | 77.67197 | 43.37133 | leaf | 9 | M. sieversii | 28 | 0 | 12KZ053 | wild | 176 | 0.0098007 |
| tuk | forest | 77.67216 | 43.37158 | fruit | 10 | M. sieversii | 27 | 0.0680969 | - | - |  | - |
| tuk | forest | 77.67203 | 43.37119 | leaf | 11 | M. sieversii | 28 | 0.0465749 | 12KZ055 | wild | 177 | 0.011822 |
| tuk | forest | 77.67193 | 43.37148 | leaf | 12 | M. sieversii | 21 | 0 | - | - |  | - |
| tuk | forest | 77.67186 | 43.37149 | leaf | 13 | M. sieversii | 21 | 0 | - | - |  | - |
| tuk | forest | 77.67187 | 43.37119 | fruit | 14 | M. sieversii | 22 | 0.0427233 | 12KZ057 | wild | 181 | 0.0239288 |
| tuk | forest | 77.67188 | 43.37158 | fruit | 15 | M. sieversii | 21 | 0.0504015 | 12KZ058 | wild | 177 | 0.0227052 |
| tuk | forest | 77.67188 | 43.37158 | leaf | 15 | M. sieversii | 21 | 0.0504015 | 12KZ059 | wild | 159 | 0.040854 |
| tuk | forest | 77.67192 | 43.37122 | fruit | 16 | M. sieversii | 23 | 0.0477525 | 12KZ060 | wild | 170 | 0.0472003 |
| tuk | forest | 77.67229 | 43.37112 | fruit | 17 | M. sieversii | 28 | 0 | 12KZ061 | wild | 180 | 0.011096 |
| tuk | forest | 77.67205 | 43.3709 | leaf | 18 | M. sieversii | 28 | 0.0342404 | 12KZ062* | hybrid | 179 | 0.4092431 |
| tuk | forest | 77.67216 | 43.37158 | leaf | 19 | M. sieversii | 25 | 0.0418651 | 12KZ064 | wild | 179 | 0.0518806 |
| tuk | forest | 77.67216 | 43.37158 | leaf | 19 | M. sieversii | 25 | 0.0418651 | 12KZ075 | wild | 178 | 0.0200468 |
| tuk | forest | 77.67229 | 43.37112 | leaf | 20 | M. sieversii | 23 | 0.0353757 | 12KZ065* | wild | 174 | 0.0205697 |
| tuk | forest | 77.67229 | 43.37112 | fruit | 20 | M. sieversii | 23 | 0.0353757 | 12KZ066* | hybrid | 178 | 0.401671 |
| tuk | forest | 77.67222 | 43.371 | leaf | 21 | M. sieversii | 28 | 0.0736853 | 12KZ067 | wild | 180 | 0.0156169 |
| tuk | forest | 77.67226 | 43.37086 | leaf | 22 | M. sieversii | 28 | 0.0736853 | 12KZ076 | wild | 177 | 0.033489 |
| tuk | forest | 77.67226 | 43.37086 | fruit | 22 | M. sieversii | 28 | 0.0736853 | 12KZ077 | wild | 175 | 0.032589 |
| tuk | forest | 77.67185 | 43.37123 | leaf | 23 | M. sieversii | 24 | 0 | - | - |  | - |
| tuk | forest | 77.67194 | 43.37121 | leaf | 24 | M. sieversii | 28 | 0.0442357 | 12KZ068 | wild | 180 | 0.0154536 |
| tuk | forest | 77.67207 | 43.37084 | leaf | 25 | M. sieversii | 28 | 0.0342404 | 12KZ069 | agricultural | 179 | 0.9744783 |
| tuk | forest | 77.67188 | 43.37115 | fruit | 26 | M. sieversii | 28 | 0.0451999 | 12KZ078 | wild | 179 | 0.0282744 |
| tuk | forest | 77.67184 | 43.37126 | fruit | 27 | M. sieversii | 28 | 0.0454225 | 12KZ074 | wild | 174 | 0.0248734 |
| tuk | forest | 77.67223 | 43.37082 | leaf | 28 | M. sieversii | 28 | 0.0736853 | 12KZ071 | wild | 177 | 0.0273656 |
| tuk | forest | 77.67233 | 43.37108 | leaf | 29 | M. sieversii | 28 | 0.057358 | 12KZ070 | agricultural | 174 | 0.9353774 |
| tuk | forest | 77.67233 | 43.37108 | leaf | 29 | M. sieversii | 28 | 0.057358 | 12KZ080* | hybrid | 175 | 0.4602008 |
| tuk | forest | 77.67214 | 43.37093 | leaf | 30 | M. sieversii | 28 | 0.0736853 | 12KZ072 | wild | 177 | 0.0322658 |
| tuk | forest | 77.67217 | 43.37164 | leaf | 31 | M. sieversii | 27 | 0 | - | - |  | - |
| tuk | forest | 77.67169 | 43.37128 | leaf | 32 | M. sieversii | 28 | 0.0492886 | - | - |  | - |
| tuk | forest | 77.67204 | 43.37128 | leaf | 33 | M. sieversii | 27 | 0.0326962 | - | - |  | - |
| tuk | forest | 77.67156 | 43.37181 | leaf | 34 | M. sieversii | 27 | 0.0423804 | - | - |  | - |
| tuk | forest | 77.67187 | 43.37159 | leaf | 35 | M. sieversii | 27 | 0.0577427 | - | - |  | - |
| tuk | forest | 77.67188 | 43.37139 | leaf | 36 | M. sieversii | 26 | 0.0330959 | - | - |  | - |
| tum | forest | 77.5832 | 43.34273 | fruit | 1 | M. sieversii | 23 | 0.0576928 | - | - |  | - |
| tum | forest | 77.58263 | 43.34264 | fruit | 2 | M. sieversii | 26 | 0 | 12KZ273 | hybrid | 176 | 0.89907 |
| tum | forest | 77.58263 | 43.34264 | fruit | 2 | M. sieversii | 26 | 0 | 12KZ345 | hybrid | 176 | 0.899147 |
| tum | forest | 77.58266 | 43.34257 | leaf | 3 | M. sieversii | 28 | 0 | 12KZ274 | wild | 167 | 0.0331265 |
| tum | forest | 77.58274 | 43.34245 | fruit | 4 | M. sieversii | 27 | 0.0370775 | 12KZ347 | wild | 179 | 0.0197248 |
| tum | forest | 77.58259 | 43.34232 | fruit | 5 | M. sieversii | 25 | 0 | 12KZ276* | hybrid | 179 | 0.4687368 |
| tum | forest | 77.58259 | 43.34232 | fruit | 5 | M. sieversii | 25 | 0 | 12KZ350 | hybrid | 177 | 0.4676094 |
| tum | forest | 77.58243 | 43.34229 | fruit | 6 | M. sieversii | 28 | 0 | 12KZ277 | wild | 179 | 0.0493067 |
| tum | forest | 77.58243 | 43.34229 | fruit | 6 | M. sieversii | 28 | 0 | 12KZ351 | wild | 176 | 0.0500835 |
| tum | forest | 77.58233 | 43.34217 | fruit | 7 | M. sieversii | 27 | 0 | 12KZ278 | wild | 179 | 0.0229511 |
| tum | forest | 77.58233 | 43.34217 | leaf | 7 | M. sieversii | 27 | 0 | 12KZ352 | hybrid | 176 | 0.8864657 |
| tum | forest | 77.58099 | 43.34187 | fruit | 8 | M. sieversii | 26 | 0 | - | - | - | - |
| tum | forest | 77.58062 | 43.34182 | fruit | 9 | M. sieversii | 24 | 0.0425736 | 12KZ280 | wild | 178 | 0.0226497 |
| tum | forest | 77.58067 | 43.34282 | leaf | 10 | M. sieversii | 23 | 0.0773487 | - | - | - | - |
| tum | forest | 77.58067 | 43.34282 | leaf | 10 | M. sieversii | 23 | 0.0773487 | 12KZ282 | hybrid | 178 | 0.6286003 |
| tum | forest | 77.58062 | 43.34198 | fruit | 11 | M. sieversii | 28 | 0 | 12KZ283 | wild | 180 | 0.0209674 |
| tum | forest | 77.58059 | 43.34196 | leaf | 12 | M. sieversii | 28 | 0 | 12KZ284 | wild | 177 | 0.0178711 |
| tum | forest | 77.58061 | 43.34198 | leaf | 13 | M. sieversii | 28 | 0 | 12KZ285 | hybrid | 179 | 0.3889651 |
| tum | forest | 77.58064 | 43.34216 | leaf | 14 | M. sieversii | 26 | 0.0464932 | 12KZ286 | wild | 180 | 0.0652164 |
| tum | forest | 77.58048 | 43.34219 | leaf | 15 | M. sieversii | 25 | 0 | 12KZ287 | wild | 174 | 0.0548881 |
| tum | forest | 77.58041 | 43.34219 | leaf | 16 | M. sieversii | 27 | 0 | - | - |  | - |
| tum | forest | 77.58031 | 43.34222 | leaf | 17 | M. sieversii | 27 | 0 | 12KZ289 | wild | 178 | 0.0141989 |
| tum | forest | 77.58026 | 43.34227 | fruit | 18 | M. sieversii | 27 | 0.0510346 | 12KZ290 | wild | 179 | 0.064826 |
| tum | forest | 77.58022 | 43.34225 | fruit | 19 | M. sieversii | 26 | 0 | 12KZ348 | wild | 178 | 0.0314614 |
| tum | forest | 77.5801 | 43.34234 | leaf | 20 | M. sieversii | 28 | 0.0433568 | 12KZ292 | hybrid | 179 | 0.1130948 |
| tum | forest | 77.58014 | 43.34234 | fruit | 21 | M. sieversii | 26 | 0.0629771 | - | - |  | - |
| tum | forest | 77.58015 | 43.34245 | leaf | 22 | M. sieversii | 26 | 0.0629771 | 12KZ294 | wild | 178 | 0.020783 |
| tum | forest | 77.58015 | 43.34251 | fruit | 23 | M. sieversii | 27 | 0.1422075 | 12KZ295 | wild | 179 | 0 |
| tum | forest | 77.58006 | 43.3427 | fruit | 24 | M. sieversii | 24 | 0.0884051 | 12KZ296 | wild | 180 | 0.0250105 |
| tum | forest | 77.58025 | 43.3428 | fruit | 25 | M. sieversii | 28 | 0.0355773 | 12KZ297 | wild | 180 | 0.0437466 |
| tum | forest | 77.58023 | 43.34279 | leaf | 26 | M. sieversii | 27 | 0.0351655 | 12KZ298 | wild | 179 | 0.0382542 |
| tum | forest | 77.58041 | 43.34273 | leaf | 27 | M. sieversii | 24 | 0 | - | - |  | - |
| tum | forest | 77.58028 | 43.3429 | leaf | 28 | M. sieversii | 27 | 0.0443097 | 12KZ300* | wild | 179 | 0.0169273 |
| tum | forest | 77.58056 | 43.34283 | fruit | 29 | M. sieversii | 27 | 0 | 12KZ301 | wild | 173 | 0.0142644 |
| tum | forest | 77.58086 | 43.3428 | fruit | 30 | M. sieversii | 25 | 0 | - | - |  | - |

**Table S2**: Description of *Venturia inaequalis* reference strains for agricultural-type and wild-type populations. These strains were sampled in 2006 in the populations Ksiev2 and Ksiev3 previously analysed in Gladieux et al. (2010).* sequenced strains number in Le Cam et al. (2019)

| Sampling location | Tree species | Strain | Fungal type | ID pop | ID strain* |
| --- | --- | --- | --- | --- | --- |
| Almaty suburbs | *M. sieversii* | 06KZ400 | agricultural | Ksiev3 | 2462 |
| Almaty suburbs | *M. sieversii* | 06KZ402 | agricultural | Ksiev3 | 2227 |
| Almaty suburbs | *M. sieversii* | 06KZ406 | agricultural | Ksiev3 | 2464 |
| Almaty suburbs | *M. sieversii* | 06KZ500 | agricultural | Ksiev3 | 2466 |
| Almaty suburbs | *M. sieversii* | 06KZ503 | agricultural | Ksiev3 | 2467 |
| Almaty suburbs | *M. sieversii* | 06KZ603 | agricultural | Ksiev3 | 2469 |
| Almaty suburbs | *M. sieversii* | 06KZ607 | agricultural | Ksiev3 | 2470 |
| Almaty suburbs | *M. sieversii* | 06KZ608 | agricultural | Ksiev3 | 2471 |
| Almaty suburbs | *M. sieversii* | 06KZ610 | agricultural | Ksiev3 | 2472 |
| Almaty suburbs | *M. sieversii* | 06KZ611 | agricultural | Ksiev3 | 2229 |
| Almaty suburbs | *M. sieversii* | 06KZ612 | agricultural | Ksiev3 | 2473 |
| Almaty suburbs | *M. sieversii* | 06KZ613 | agricultural | Ksiev3 | 2474 |
| Almaty suburbs | *M. sieversii* | 06KZ617 | agricultural | Ksiev3 | 2476 |
| Almaty suburbs | *M. sieversii* | 06KZ704 | agricultural | Ksiev3 | 2230 |
| Almaty suburbs | *M. sieversii* | 06KZ709 | agricultural | Ksiev3 | 2232 |
| Kourznetsov | *M. sieversii* | 06KZ304 | agricultural | Ksiev2 | 2448 |
| Kourznetsov | *M. sieversii* | 06KZ308 | agricultural | Ksiev2 | 2449 |
| Kourznetsov | *M. sieversii* | 06KZ314 | agricultural | Ksiev2 | 2452 |
| Kourznetsov | *M. sieversii* | 06KZ309 | wild | Ksiev2 | 2450 |
| Kourznetsov | *M. sieversii* | 06KZ312a | wild | Ksiev2 | 2451 |
| Kourznetsov | *M. sieversii* | 06KZ316 | wild | Ksiev2 | 2453 |
| Kourznetsov | *M. sieversii* | 06KZ317 | wild | Ksiev2 | 2454 |
| Kourznetsov | *M. sieversii* | 06KZ318b | wild | Ksiev2 | 2223 |
| Kourznetsov | *M. sieversii* | 06KZ320b | wild | Ksiev2 | 2455 |
| Kourznetsov | *M. sieversii* | 06KZ321 | wild | Ksiev2 | 2456 |
| Kourznetsov | *M. sieversii* | 06KZ324 | wild | Ksiev2 | 2457 |
| Kourznetsov | *M. sieversii* | 06KZ328 | wild | Ksiev2 | 2459 |
| Kourznetsov | *M. sieversii* | 06KZ330 | wild | Ksiev2 | 2460 |
| Kourznetsov | *M. sieversii* | 06KZ332 | wild | Ksiev2 | 2461 |
| Kournetsov | *M. sieversii* | 06KZ409 | wild | Ksiev2 | 2465 |

**Table S3:** Information on the 192 SNPs used for *Venturia inaequalis* genotyping: name, location (scaffold and physical position in bp) on the reference genome EU-B04 (NCBI Accession number: ASM368922v1), the alleles, and the F_ST_ estimate between wild and agricultural populations are given. The 181 SNPs finally used in this paper are indicated in bold and italic.

| SNP Name | Scaffold on EUB04 | Position (bp) | Alleles | F_ST_ |
| --- | --- | --- | --- | --- |
| ***V_018220_1331*** | VeinUTG001 | 839254 | C/T | 0.8028 |
| ***V_018840_0909*** | VeinUTG001 | 1039813 | C/T | 0.8723 |
| ***V_019170_0355*** | VeinUTG001 | 1117770 | G/C | 0.7355 |
| ***V_029490_0190*** | VeinUTG001 | 1541049 | A/G | 0.9351 |
| ***V_126720_0104*** | VeinUTG001 | 1808566 | A/G | 0.7470 |
| ***V_109090_0141*** | VeinUTG001 | 1954918 | C/T | 1.0000 |
| ***V_041540_1577*** | VeinUTG001 | 2247693 | A/T | 0.7470 |
| ***V_035660_0936*** | VeinUTG001 | 3031026 | A/T | 0.8694 |
| ***V_105310_1263*** | VeinUTG001 | 3265132 | C/T | 0.8694 |
| ***V_079730_1403*** | VeinUTG001 | 3995960 | A/G | 0.7355 |
| ***V_023320_2232*** | VeinUTG001 | 4236708 | G/C | 0.9358 |
| ***V_111790_0550*** | VeinUTG001 | 4439887 | G/C | 0.8093 |
| ***V_032790_0216*** | VeinUTG001 | 4703816 | A/G | 0.8630 |
| ***V_066160_0089*** | VeinUTG002 | 765559 | C/T | 0.7918 |
| ***V_110640_0020*** | VeinUTG002 | 1561889 | C/T | 1.0000 |
| ***V_107940_0512*** | VeinUTG002 | 1738342 | C/T | 0.7470 |
| ***V_099640_0224*** | VeinUTG002 | 1838866 | A/G | 0.8723 |
| ***V_052850_0506*** | VeinUTG002 | 2018870 | G/T | 1.0000 |
| ***V_118140_0086*** | VeinUTG002 | 2498819 | C/T | 0.8028 |
| ***V_085920_0461*** | VeinUTG002 | 2642234 | C/T | 0.7470 |
| ***V_046550_2040*** | VeinUTG002 | 3194538 | A/C | 0.8093 |
| ***V_113090_0515*** | VeinUTG002 | 3466914 | C/T | 0.7470 |
| ***V_097550_0704*** | VeinUTG002 | 3977279 | A/C | 1.0000 |
| ***V_100730_0065*** | VeinUTG002 | 4147318 | A/G | 1.0000 |
| ***V_116690_0737*** | VeinUTG003 | 167073 | C/T | 0.9358 |
| ***V_079380_0627*** | VeinUTG003 | 564968 | G/T | 1.0000 |
| ***V_054000_0204*** | VeinUTG003 | 1037767 | A/G | 1.0000 |
| ***V_100130_0367*** | VeinUTG003 | 1465410 | A/T | 0.8694 |
| ***V_067250_0582*** | VeinUTG003 | 1844911 | A/T | 0.9351 |
| ***V_116430_0046*** | VeinUTG003 | 2149034 | A/G | 0.8571 |
| ***V_081690_0319*** | VeinUTG003 | 2409684 | A/G | 1.0000 |
| V_120330_1617 | VeinUTG003 | 3111881 | A/G | 0.8723 |
| ***V_057600_0725*** | VeinUTG003 | 3317502 | C/T | 0.6853 |
| ***V_056430_1439*** | VeinUTG003 | 3742624 | G/T | 0.7470 |
| ***V_074810_0013*** | VeinUTG003 | 4069197 | C/T | 1.0000 |
| ***V_111300_0184*** | VeinUTG004 | 366077 | G/T | 1.0000 |
| ***V_110380_1794*** | VeinUTG004 | 514643 | A/G | 1.0000 |
| ***V_037930_3318*** | VeinUTG004 | 1018024 | C/T | 0.5637 |
| ***V_036860_0360*** | VeinUTG004 | 1345820 | A/G | 0.8093 |
| ***V_096550_1125*** | VeinUTG004 | 2272628 | A/C | 1.0000 |
| ***V_053560_0224*** | VeinUTG004 | 2674971 | C/T | 0.7470 |
| ***V_092240_0011*** | VeinUTG004 | 3775788 | A/C | 1.0000 |
| ***V_112200_0034*** | VeinUTG004 | 3921677 | A/G | 0.9358 |
| ***V_102170_0137*** | VeinUTG005 | 458201 | A/C | 0.6242 |
| ***V_087620_0047*** | VeinUTG005 | 634777 | C/T | 0.9351 |
| ***V_089860_0444*** | VeinUTG005 | 1137669 | C/T | 0.7918 |
| ***V_090230_0543*** | VeinUTG005 | 1237890 | C/T | 1.0000 |
| ***V_130920_0164*** | VeinUTG005 | 1298244 | A/T | 0.8093 |
| ***V_117750_1887*** | VeinUTG005 | 1378629 | A/T | 0.5470 |
| ***V_060730_0222*** | VeinUTG005 | 1845447 | C/T | 0.8093 |
| ***V_019860_0666*** | VeinUTG005 | 1949024 | C/T | 0.8723 |
| ***V_020850_0531*** | VeinUTG005 | 2258475 | A/G | 0.6853 |
| ***V_022570_0172*** | VeinUTG005 | 2811988 | A/G | 0.7918 |
| ***V_022940_0143*** | VeinUTG005 | 2891444 | G/T | 0.9358 |
| ***V_008320_0134*** | VeinUTG005 | 3107923 | A/C | 0.9358 |
| ***V_008780_0414*** | VeinUTG005 | 3232243 | A/G | 1.0000 |
| ***V_098950_0212*** | VeinUTG005 | 3429229 | G/C | 1.0000 |
| ***V_128170_0756*** | VeinUTG005 | 3766438 | A/C | 0.9358 |
| ***V_123170_2529*** | VeinUTG005 | 3924343 | A/G | 0.8723 |
| ***V_073830_0552*** | VeinUTG006 | 102410 | G/T | 0.7470 |
| ***V_141550_0483*** | VeinUTG006 | 343249 | C/T | 0.7355 |
| ***V_078180_0144*** | VeinUTG006 | 535658 | C/T | 1.0000 |
| ***V_015030_0264*** | VeinUTG006 | 730418 | C/T | 0.8723 |
| ***V_014480_0440*** | VeinUTG006 | 888941 | C/T | 0.9351 |
| V_013530_0158 | VeinUTG006 | 1127706 | C/T | 0.8914 |
| ***V_134340_1382*** | VeinUTG006 | 1291066 | A/G | 0.8694 |
| ***V_102910_0947*** | VeinUTG006 | 1651347 | C/T | 1.0000 |
| ***V_090440_0036*** | VeinUTG006 | 1801383 | C/T | 1.0000 |
| ***V_065050_1034*** | VeinUTG006 | 2055794 | C/T | 0.9351 |
| ***V_111050_1271*** | VeinUTG006 | 2971000 | C/T | 0.7470 |
| ***V_062600_0140*** | VeinUTG006 | 3049397 | A/G | 0.8093 |
| ***V_113600_0900*** | VeinUTG006 | 3854925 | A/T | 1.0000 |
| V_095590_1275 | VeinUTG008 | 484919 | A/G | 0.3274 |
| ***V_123720_1795*** | VeinUTG008 | 583358 | G/T | 0.8694 |
| ***V_059540_0717*** | VeinUTG008 | 1074360 | C/T | 0.7216 |
| ***V_077220_2072*** | VeinUTG008 | 1261806 | C/T | 0.9351 |
| ***V_077590_0210*** | VeinUTG008 | 1347572 | A/C | 1.0000 |
| ***V_010740_0551*** | VeinUTG008 | 1641781 | A/C | 0.7149 |
| ***V_009670_0207*** | VeinUTG008 | 1942030 | C/T | 1.0000 |
| ***V_009480_0124*** | VeinUTG008 | 2009829 | A/G | 0.8093 |
| ***V_137400_1007*** | VeinUTG008 | 2192869 | A/G | 0.9351 |
| ***V_131300_0261*** | VeinUTG008 | 2393042 | A/G | 0.3274 |
| V_107140_0544 | VeinUTG008 | 2478815 | A/G | 0.2561 |
| ***V_140080_0973*** | VeinUTG008 | 2770084 | C/T | 1.0000 |
| ***V_129680_1329*** | VeinUTG008 | 3204417 | A/T | 0.7918 |
| ***V_103130_0267*** | VeinUTG009 | 224647 | A/C | 0.9351 |
| ***V_088800_0465*** | VeinUTG009 | 312067 | C/T | 0.8630 |
| ***V_088850_1431*** | VeinUTG009 | 324021 | A/T | 1.0000 |
| ***V_003980_1058*** | VeinUTG009 | 712661 | A/G | 1.0000 |
| ***V_003160_1629*** | VeinUTG009 | 975046 | C/T | 0.9358 |
| ***V_001910_1422*** | VeinUTG009 | 1286243 | C/T | 0.5169 |
| ***V_000760_0258*** | VeinUTG009 | 1563963 | G/T | 0.7918 |
| ***V_133290_0122*** | VeinUTG009 | 1853322 | A/G | 0.9358 |
| ***V_017940_0347*** | VeinUTG009 | 2030072 | C/T | 0.7470 |
| V_016950_0318 | VeinUTG009 | 2498049 | G/C | 0.5038 |
| ***V_112740_0817*** | VeinUTG009 | 2796178 | A/G | 0.9351 |
| ***V_100300_2012*** | VeinUTG010 | 10417 | A/G | 0.8093 |
| ***V_055490_0575*** | VeinUTG010 | 451375 | A/G | 0.8630 |
| ***V_044450_0210*** | VeinUTG010 | 653247 | C/T | 1.0000 |
| ***V_102610_1164*** | VeinUTG010 | 948131 | A/C | 1.0000 |
| ***V_124510_1254*** | VeinUTG010 | 1086716 | A/C | 0.5038 |
| ***V_026720_1062*** | VeinUTG010 | 1464856 | A/T | 0.9358 |
| ***V_084190_0186*** | VeinUTG010 | 1954158 | C/T | 1.0000 |
| ***V_106710_0135*** | VeinUTG010 | 2130383 | C/T | 1.0000 |
| ***V_115300_0879*** | VeinUTG010 | 2373680 | A/C | 0.6242 |
| ***V_115680_2534*** | VeinUTG010 | 2727345 | A/G | 0.8723 |
| ***V_108180_0131*** | VeinUTG011 | 314465 | A/G | 0.9351 |
| ***V_076460_0566*** | VeinUTG011 | 688867 | A/G | 0.5637 |
| ***V_121930_1112*** | VeinUTG011 | 809888 | G/T | 0.7470 |
| ***V_117670_0236*** | VeinUTG011 | 1160812 | A/C | 0.9351 |
| ***V_058530_1413*** | VeinUTG011 | 2213367 | C/T | 1.0000 |
| ***V_092800_0959*** | VeinUTG011 | 2524034 | A/G | 0.8093 |
| ***V_106060_0500*** | VeinUTG011 | 2681070 | A/T | 0.8694 |
| ***V_062170_0122*** | VeinUTG012 | 358671 | C/T | 0.9351 |
| ***V_031690_0750*** | VeinUTG012 | 672423 | A/G | 0.8630 |
| ***V_118540_0041*** | VeinUTG012 | 1095999 | C/T | 0.9358 |
| ***V_093830_0965*** | VeinUTG012 | 1386051 | C/T | 0.5038 |
| ***V_073430_0035*** | VeinUTG012 | 1632290 | C/T | 0.9358 |
| ***V_094490_0565*** | VeinUTG012 | 1992493 | A/G | 0.9358 |
| ***V_117350_0545*** | VeinUTG013 | 341065 | A/G | 0.7692 |
| ***V_024350_0126*** | VeinUTG013 | 632459 | C/T | 0.8093 |
| ***V_063460_1376*** | VeinUTG013 | 2024135 | A/C | 0.2117 |
| ***V_061560_0428*** | VeinUTG013 | 2356257 | C/T | 0.8694 |
| ***V_075040_0251*** | VeinUTG014 | 579982 | C/T | 0.9358 |
| ***V_050770_0063*** | VeinUTG014 | 934352 | G/C | 1.0000 |
| ***V_104100_0053*** | VeinUTG014 | 1341421 | A/G | 1.0000 |
| ***V_120670_1164*** | VeinUTG014 | 1562110 | G/T | 0.8028 |
| ***V_101260_0476*** | VeinUTG014 | 1664063 | G/C | 1.0000 |
| ***V_103700_0044*** | VeinUTG014 | 2154755 | A/G | 0.8630 |
| ***V_070750_1596*** | VeinUTG015 | 302330 | A/G | 0.8028 |
| ***V_069780_1161*** | VeinUTG015 | 778380 | G/C | 0.9351 |
| ***V_081980_2018*** | VeinUTG015 | 1204853 | A/G | 0.5637 |
| ***V_045450_0080*** | VeinUTG015 | 1774969 | A/G | 0.9064 |
| ***V_004980_1116*** | VeinUTG016 | 141273 | C/T | 1.0000 |
| ***V_005510_0793*** | VeinUTG016 | 296120 | A/G | 0.9351 |
| ***V_006120_1278*** | VeinUTG016 | 469371 | C/T | 1.0000 |
| ***V_006370_0207*** | VeinUTG016 | 531479 | C/T | 1.0000 |
| ***V_007020_0140*** | VeinUTG016 | 794963 | A/G | 0.8694 |
| ***V_129790_0400*** | VeinUTG016 | 879607 | C/T | 1.0000 |
| ***V_129830_0447*** | VeinUTG016 | 927418 | C/T | 0.9358 |
| ***V_096110_0527*** | VeinUTG016 | 1155149 | A/C | 0.5590 |
| ***V_040250_0011*** | VeinUTG016 | 1886600 | A/G | 1.0000 |
| ***V_033700_1050*** | VeinUTG017 | 100528 | A/G | 0.8630 |
| ***V_110060_0035*** | VeinUTG017 | 611756 | C/T | 1.0000 |
| ***V_007370_0102*** | VeinUTG017 | 1931433 | A/C | 1.0000 |
| ***V_039130_0455*** | VeinUTG018 | 937349 | C/T | 0.9351 |
| V_051430_0257 | VeinUTG018 | 1391326 | A/G | 0.8723 |
| ***V_114790_0032*** | VeinUTG018 | 1643150 | A/C | 0.8723 |
| ***V_091190_0926*** | VeinUTG019 | 656797 | A/T | 0.8093 |
| V_069210_1034 | VeinUTG019 | 873830 | A/G | 0.8630 |
| ***V_121510_1421*** | VeinUTG019 | 1190047 | C/T | 0.8723 |
| ***V_120100_0992*** | VeinUTG019 | 1305566 | C/T | 1.0000 |
| ***V_118700_0239*** | VeinUTG019 | 1623334 | C/T | 0.9351 |
| ***V_102000_0268*** | VeinUTG020 | 166969 | A/G | 0.8723 |
| V_141260_0912 | VeinUTG020 | 215237 | A/G | 0.8723 |
| ***V_012620_0888*** | VeinUTG020 | 400156 | C/T | 0.8723 |
| ***V_011270_0106*** | VeinUTG020 | 727448 | A/C | 0.3857 |
| ***V_098010_1257*** | VeinUTG020 | 934007 | A/C | 0.8093 |
| ***V_093290_0426*** | VeinUTG020 | 1104985 | A/G | 0.8723 |
| ***V_100630_0117*** | VeinUTG020 | 1369974 | A/T | 0.7893 |
| ***V_134230_0238*** | VeinUTG020 | 1525411 | A/C | 0.5169 |
| ***V_050000_1196*** | VeinUTG021 | 205566 | G/T | 1.0000 |
| ***V_095220_0593*** | VeinUTG021 | 733672 | C/T | 0.5841 |
| ***V_043620_0935*** | VeinUTG021 | 1207518 | G/C | 0.7216 |
| ***V_089570_0572*** | VeinUTG021 | 1551622 | A/G | 0.7149 |
| ***V_119510_0951*** | VeinUTG022 | 59072 | A/T | 0.9351 |
| ***V_107330_1521*** | VeinUTG022 | 285877 | C/T | 0.8630 |
| ***V_113470_0591*** | VeinUTG022 | 605018 | G/T | 0.7470 |
| ***V_015520_0324*** | VeinUTG022 | 995272 | C/T | 1.0000 |
| ***V_016610_0144*** | VeinUTG022 | 1330750 | C/T | 0.8093 |
| V_101310_1396 | VeinUTG023 | 53235 | A/C | 0.8093 |
| ***V_070470_0198*** | VeinUTG023 | 328925 | A/G | 0.8723 |
| ***V_080900_0945*** | VeinUTG023 | 663789 | A/C | 0.4609 |
| ***V_087410_1022*** | VeinUTG023 | 1393223 | G/C | 0.8694 |
| ***V_088520_0150*** | VeinUTG024 | 18349 | A/T | 0.7464 |
| ***V_135930_0687*** | VeinUTG024 | 456348 | A/G | 0.9358 |
| V_042990_0437 | VeinUTG024 | 823804 | A/C | 0.7893 |
| ***V_028380_1199*** | VeinUTG024 | 1067196 | A/C | 0.5481 |
| ***V_066400_1022*** | VeinUTG025 | 589959 | C/T | 0.9351 |
| ***V_098930_0257*** | VeinUTG025 | 966469 | A/T | 0.8408 |
| ***V_082830_0977*** | VeinUTG025 | 1127819 | C/T | 0.8028 |
| ***V_064670_0035*** | VeinUTG026 | 76001 | C/T | 0.8694 |
| ***V_068610_0191*** | VeinUTG026 | 489046 | C/T | 0.5169 |
| ***V_071360_0062*** | VeinUTG026 | 629492 | C/T | 0.9351 |
| ***V_098350_2060*** | VeinUTG026 | 1034092 | C/T | 0.8093 |
| ***V_116040_0960*** | VeinUTG027 | 144346 | A/G | 0.9358 |
| ***V_098380_0256*** | VeinUTG027 | 342149 | C/T | 0.7355 |
| ***V_114120_1789*** | VeinUTG027 | 696671 | A/G | 1.0000 |
| ***V_049020_0011*** | VeinUTG028 | 237295 | C/T | 0.8723 |
| ***V_113930_2603*** | VeinUTG028 | 464425 | C/T | 0.8093 |
| ***V_114950_1466*** | VeinUTG028 | 726541 | G/T | 0.8723 |
| V_131300_0261 | VeinUTG129 | 748 | A/G | 1.0000 |

**Table S4:** Expected genetic heterozygosity (*H_e_*) taken as measures of diversity, per population, for apple trees and *Venturia inaequalis.*

| Sampling sites | Apple tree *H_e_* | *V. inaequalis* *H_e_* |
| --- | --- | --- |
| kot | 0.8023 | 0.4869 |
| tag | 0.7476 | 0.4729 |
| tar | 0.7446 | 0.4558 |
| lac | 0.7809 | 0.4366 |
| gec | 0.7086 | 0.4862 |
| esi_o | 0.7389 | 0.1778 |
| esi_f | 0.7926 | 0.4679 |
| tum | 0.7179 | 0.3041 |
| tuk | 0.703 | 0.2438 |

**Table S5:** Pairwise *F_ST_* estimates between apple trees sampling sites. Significant values (*F_ST_*>0) at p-value<0.001 are indicated in bold.

|  | kot | tag | tar | lac | gec | esi_o | esi_f | tum |
| --- | --- | --- | --- | --- | --- | --- | --- | --- |
| tag | **0.005** |  |  |  |  |  |  |  |
| tar | **0.005** | **0.012** |  |  |  |  |  |  |
| lac | 0.000 | **0.016** | **0.012** |  |  |  |  |  |
| gec | **0.023** | **0.020** | **0.026** | **0.018** |  |  |  |  |
| esi_o | **0.134** | **0.160** | **0.162** | **0.129** | **0.198** |  |  |  |
| esi_f | 0.000 | **0.015** | **0.013** | **0.002** | **0.025** | **0.138** |  |  |
| tum | **0.020** | **0.022** | **0.012** | **0.016** | **0.020** | **0.189** | **0.017** |  |
| tuk | **0.031** | **0.032** | **0.033** | **0.027** | **0.036** | **0.207** | **0.030** | **0.020** |

**Table S6:** Pairwise *F_ST_* estimates between *Venturia inaequalis* sampling sites. Significant values (F_ST_ >0) at p-value<0.001 are indicated in bold.

|  | kot | tag | tar | lac | gec | esi_o | esi_f | tum |
| --- | --- | --- | --- | --- | --- | --- | --- | --- |
| tag | 0.000 |  |  |  |  |  |  |  |
| tar | 0.000 |  |  |  |  |  |  |  |
| lac | 0.004 | 0.000 | 0.000 |  |  |  |  |  |
| gec | 0.000 | 0.000 | 0.000 | 0.013 |  |  |  |  |
| esi_o | **0.370** | **0.435** | **0.475** | **0.494** | **0.357** |  |  |  |
| esi_f | **0.029** | **0.060** | **0.089** | **0.115** | 0.016 | **0.197** |  |  |
| tum | **0.117** | **0.073** | **0.050** | **0.032** | **0.135** | **0.662** | **0.281** |  |
| tuk | **0.168** | **0.123** | **0.094** | **0.070** | **0.192** | **0.721** | **0.338** | 0.000 |
